# Spatial biology of Ising-like synthetic genetic networks

**DOI:** 10.1101/2023.05.10.540292

**Authors:** Kevin Simpson, Alfredo L’Homme, Juan Keymer, Fernán Federici

## Abstract

**Background:** Understanding how spatial patterns of gene expression emerge from the interaction of individual gene networks is a fundamental challenge in biology. Developing a synthetic experimental system with a common theoretical framework that captures the emergence of short- and long-range spatial correlations (and anti-correlations) from interacting gene networks could serve to uncover generic scaling properties of these ubiquitous phenomena.

**Results:** Here, we combine synthetic biology, statistical mechanics models and computational simulations to study the spatial behavior of synthetic gene networks (SGNs) in *Escherichia coli* quasi-2D colonies growing on hard agar. Guided by the combined mechanisms of the contact process lattice simulation and two-dimensional Ising model (CPIM), we describe the spatial behavior of bi-stable and chemically-coupled SGNs that self-organize into patterns of long-range correlations with power-law scaling or short-range anti-correlations. These patterns, resembling ferromagnetic and anti-ferromagnetic configurations of the Ising model near critical points, maintain their scaling properties upon changes in growth rate and cell shape.

**Conclusions:** Our findings shed light on the spatial biology of coupled and bistable gene networks in growing cell populations. This emergent spatial behavior could provide insights into the study and engineering of self-organizing gene patterns in eukaryotic tissues and bacterial consortia.

## 1 Background

The emergence of spatially-correlated structures is a phenomenon that pervades biology from molecular to ecological scales (e.g. Noble *et al* (2018); Bray and Duke (2004); Duke *et al* (2001); Turing (1990)). An emblematic case of research is the spatial correlations of gene expression in eukaryotic tissues and microbial communities, which can occur at short-range or at the scale of the whole population (e.g.van Vliet *et al* (2018); Rosenthal *et al* (2018); Kim *et al* (2019); van Gestel *et al* (2021)). For instance, negative spatial correlations can emerge during eukaryotic cell differentiation (e.g.Collier et al (1996)) and metabolic crossfeeding in microbial systems (e.g.Dal Co et al (2020); Cole et al (2015); Lovley (2017)), whereas positive gene spatial correlations can be observed during the synchronization of growth and resource sharing in bacterial populations (e.g. Liu *et al* (2017, 2015); van Vliet *et al* (2022)). These spatial patterns are shaped from the bottom-up through mechanisms that differ widely across different organisms. Developing a common experimental system and theoretical framework that captures the formation of shortand long-range spatial correlations (and anti-correlations) from interacting gene networks could serve to uncover generic mechanisms and scaling properties of these ubiquitous phenomena. The required theoretical framework should embody the emergence of global correlations from the collective behavior of interacting gene networks in space. The Ising model from statistical mechanics is suitable for this task since it provides a mathematical machinery, amenable to simple numerical simulations and exact analysis, able to address the collective behaviour of spatially-interacting particles. Originally formulated to understand the loss of magnetism in ferromagnetic materials as the temperature increases, this model has been useful to study second order phase transitions and critical phenomena Ising (1925); Kobe (1997); Solé (2011). Different studies have demonstrated the applicability of the Ising model to investigate the spatial organization of biological processes at molecular, cellular and ecological scales. It has been used to explain the propagation of allosteric states in large multi-protein complexes Bray and Duke (2004); Duke et al (2001) as well as the emergence of long-range synchronization in ecological systems Noble *et al* (2015, 2018). At a cellular level, the Ising model has been used to study follicle alignment during mammalian hair patterning Wang et al (2006), ferromagnetic and anti-ferromagnetic correlations in lattices of hydrodynamically coupled bacterial vortices Wioland *et al* (2016). An interesting approach, by Weber and Buceta (2016), studies second order phase transitions in simulations of bacterial cell ensembles carrying coupled toggle switches Weber and Buceta (2016). Although their theoretical ensembles lack spatially-explicit structure, their numerical simulations suggest that gene regulatory networks constructed of toggle switches interconnected with quorum sensing signals can exhibit spontaneous symmetry breaking. This work suggests a minimal framework applying the Ising Model and critical phenomena to understand phase transitions in groups of cells that exhibit alternative phenotypes. Extending this approach to apply the Ising model to the study of gene spatial correlations in natural cell populations could be challenging since these systems are embedded in complex physiological contexts affected by unknown components and unforeseen interactions. Alternatively, this phenomenon could be studied in synthetic gene networks (SGNs) that embody the essential features of the Ising model. The use of SGNs as test-beds to challenge biological theory has gained popularity since it provides more experimental control and analytical power Gardner et al (2000); Elowitz and Leibler (2000); Mukherji and van Oudenaarden (2009); Elowitz and Lim (2010); Davies (2017). The use of efficient DNA fabrication methods, well-characterized components and mathematical modeling has enabled the engineering SGNs of unprecedented scale and predictability (e.g. Nielsen et al (2016)). This approach has enabled the engineering of biological patterns, a new frontier of interdisciplinary research that employs minimal and reconfigurable SGNs Luo et al (2019); Schaerli et al (2014); Basu et al (2005); Payne et al (2013); Toda et al (2018); Ebrahimkhani and Ebisuya (2019). The use of controllable SGNs embodying Ising model rules could be instrumental for defining a common theoretical ground for the spatial biology of gene networks.

Here, we apply a theoretical framework based on the Ising model to study how spatial correlations emerge from chemically-coupled, bistable SGNs in *Escherichia coli* growing on hard agar as quasi-2D colonies. By analogy with the two-state interacting particles of the model, we construct synthetic toggle switches Gardner et al (2000) whose states are chemically-coupled by quorum sensing signaling Grant et al (2016). These SGNs self-organize in long-range spatial correlations and fractal patterns reminiscent of ferromagnetic systems of the Ising model. Inverting the response to the coupling signals, on the other hand, creates negative correlations similar to anti-ferromagnetic configurations, demonstrating correspondence between SGNs and the model.

## 2 Results

### 2.1 A two-dimensional Ising model in growing cell populations

To study how long-range gene correlations arise from diffusion-limited chemical coupling between gene networks, we employed the Ising model with two-state particles arranged on a two-dimensional lattice (Fig. 1a). To implement it in a growing population of cells, we generated a lattice model that combines the Contact Process Harris (1974), representing cell population dynamics, with the two-dimensional Ising model using the Metropolis algorithm Metropolis et al (1953). This model, hereafter named CPIM (for “Contact Process Ising Model”), consists of an interacting particle system that follows colonization, extinction, and differentiation dynamics of particles on a two-dimensional lattice ℒ of *N* sites (Fig. 1b, Additional file 2, Fig. S16, Supplementary Movie 1). As in the Ising model, cells are fixed in their position and can only interact with their nearest four neighbors with interaction energy ℋ, which in the absence of external perturbations is determined by the Hamiltonian:

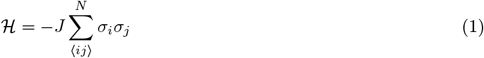

where *σ* corresponds to the state of differentiated cells (magenta or green, +1 or −1), *ij* indicates that the sum is only between neighboring pairs of cells (only short-range interactions are allowed), and *J* represents the contribution to free energy made by the coupling interaction between these pairs of cells. We assumed that *J* = +1 for ferromagnetic systems and *J* = −1 for anti-ferromagnetic systems. This implies that the interaction energy between neighboring cells is minimized when they have the same state in a ferromagnetic system, or the opposite state in an anti-ferromagnetic system (Eq. 1). In the Ising model, the probability *P* (in equilibrium) of a certain spatial configuration of spins *x* is defined by 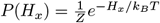, where *k*_*B*_is the Boltzmann constant, *Z* is the partition function 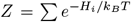 (sum over all possible configurations), and *T* is the absolute temperature of the system. Together with Eq. 1, these equations show the contribution of *J* and *T* to the probability distribution of spatial configurations: for a given value of *J*, varying *T* determines the transition between ordered and disordered configurations of the system (Fig. 1a). Since we are interested in studying the emergence of gene correlations due to coupling between gene networks, in CPIM, *T* represents a parameter that regulates coupling in the system: a small value of *T* allows for a strong coupling, while a large value represents a weak coupling.

**Figure 1:**
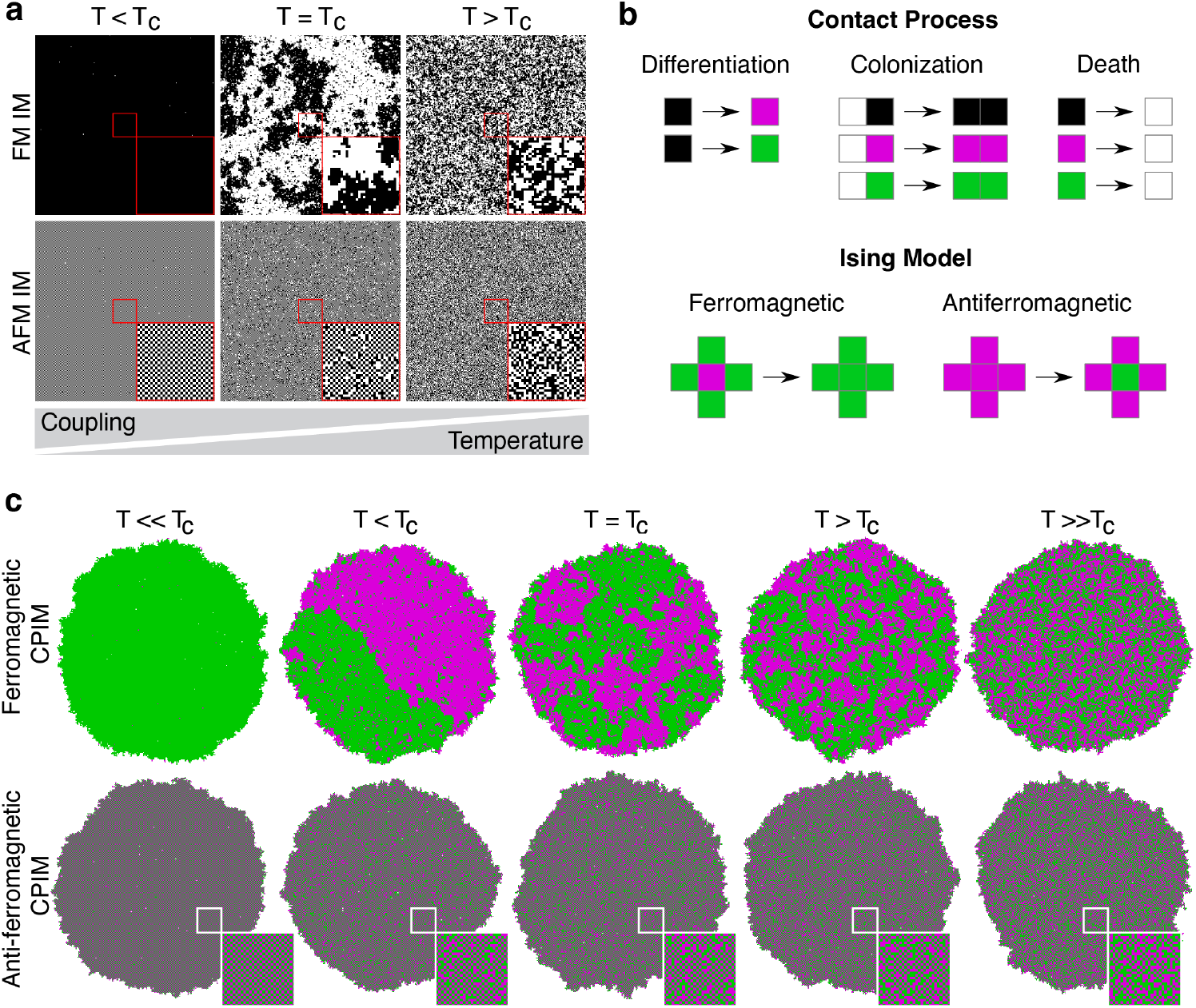
Ising-like interactions in a growing population of cells. (a) Numerical simulations of the two-dimensional ferromagnetic (FM IM) and anti-ferromagnetic (AFM IM) Ising model on a 250×250 lattice at different temperatures *T* relative to the critical temperature *T*_*c*_. White and black squares represent the spin orientations *σ* = ±1. Insets correspond to a magnification of a 30×30 square in the center of the images. (b) Basic rules that define CPIM simulations: Contact process lattice reactions of colonization, differentiation and death processes (top); and Ising-like cellular state change mechanisms (bottom). Each site of this lattice can be in one of four states *S* = ∅∗, +1,−1, which represent vacant locations (∅, white squares), locations occupied by undifferentiated cells (∗, black squares), and locations occupied by differentiated cells in red (+1, magenta squares) or green (−1, green squares) state. (c) CPIM numerical simulations of growing ferromagnetic and anti-ferromagnetic cell populations at different values of the control parameter *T* relative to the critical value *T*_*c*_. In the CPIM simulations, *T* represents a parameter that determines the strength of coupling between cells. Insets show a magnification of the square in the center of anti-ferromagnetic colonies showing a detail of the checkerboard-like pattern.

The net magnetization of a population (*M*), which represents the order parameter of the system, is determined by the degree of alignment of the cells, and is given by *M* = £ _*i*_ *s*_*i*_. The mean magnetization per site (*M/N*) calculated for ferromagnetic populations simulated with CPIM (Additional file 1, Fig. S1A) showed good agreement with the behavior observed in the Ising model Ibarra-García-Padilla et al (2016). Fig. 1c shows the different cellular state configurations that emerge in simulated populations depending on the strength of the coupling (the value of *T*) and the type of interaction (ferromagnetic or anti-ferromagnetic). In a ferromagnetic population (Fig. 1c top), a strong interaction between cells (*T < T*_*c*_) favors the alignment of states, leading to the emergence of large homogeneous patches of the same state. A further increase in the coupling (*T << T*_*c*_) leads to colonies with cells in only one state. On the other hand, when the coupling between cells is weak (*T > T*_*c*_), cells freely change their states regardless of the state of their neighbors, leading to a colony with a noise-like appearance. Near a critical value of coupling (*T* = *T*_*c*_), colonies spontaneously self-organize into patterns that resemble the long-range correlations and power-law decaying fractal objects described by universality class exponents of the Ising model at the phase transition. At this critical point, the autocorrelation function of the simulated populations follows a power-law decay given by 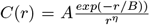 (Additional file 1, Fig. S1C), with a critical exponent of the autocorrelation function *η* = 0.2518, consistent with the value reported for the Ising model at the critical temperature (*η* = 0.250) Ibarra-García-Padilla *et al* (2016). This behavior close to the critical transition is particularly relevant since it links the short-range coupling between cellular states to the generation of macroscopic long-range correlations.

In anti-ferromagnetic populations, an ordered configuration emerges under strong interactions between cells and a disordered configuration is observed when the interaction is weak (Fig. 1c bottom). However, opposite states between neighboring cells are favored in the ordered configuration, resulting in the emergence of a checkerboard-like pattern of cellular states (a red cell surrounded by four green cells and vice versa). Near the critical value, colonies are composed of patches of checkerboard-like patterns separated by disordered regions.

These CPIM simulations showed that growing cell populations with ferromagnetic and antiferromagnetic Ising-like interactions give rise to short and long-range correlations around critical points. This suggests that SGNs with two states coupled through chemical signals that capture ferromagnetic and antiferromagnetic interactions should lead to spatial patterns of positive or negative correlations, respectively.

### Synthetic gene networks with spin-like behaviour

We constructed synthetic gene circuits capturing Ising mechanisms, i.e. two states and coupling interactions (Fig. 2). We named these systems ferromagnetic or anti-ferromagnetic depending on whether they promote the same or opposite state in their neighbors, respectively (Fig. 2a). Each system is composed of three main functions: *i*) a state reporter, responsible for the synthesis of a red fluorescent protein (mCherry2, hereafter called “RFP” for simplicity) or a green fluorescent protein (sfGFP, hereafter called “GFP” for simplicity) to report the state of SGNs, *ii*) a switch module, composed of two repressors (LacI and TetR) that repress the expression of each other, allowing cells to adopt only one of the two possible states at a time, and *iii*) a coupling module, which allows the production of one of the two coupling signals for each state (3-oxo-C6-homoserine lactone (C6HSL), synthesized by LuxI from *Vibrio fischeri*, or 3-oxo-C12homoserine lactone (C12HSL), synthesized by LasI from *Pseudomonas aeruginosa*). These functions are contained in two vectors: the reporter vector and the ferromagnetic or anti-ferromagnetic vector (Fig. 2b). The expression of genes encoding the repressors, red/green fluorescent protein, and C6HSL and C12HSL biosynthetic enzymes are under the control of two inducible/repressible promoters: the pLuxpTet promoter (induced by C6HSL and repressed by TetR) and the pLaspLac promoter (induced by C12HSL and repressed by LacI). To gain orthogonality, we used the pLux76 and pLas81 versions of the pLux and pLas promoters, respectively Grant et al (2016) (hereafter named pLux76pTet and pLas81pLac). Thus, the states of these SGNs are determined by the coupling molecules and their ferromagnetic or anti-ferromagnetic configurations. In the ferromagnetic vector, cells synthesize the same coupling molecule they sense, inducing the same state in neighboring cells. Conversely, in the anti-ferromagnetic vector, cells synthesize the opposite coupling signal they sense, inducing the opposite state in neighboring cells. In both systems, the *LacI* and *TetR* repressors are under the pLux76pTet and pLas81pLac promoters, respectively, ensuring that the production of one coupling signal is accompanied by the repression of the other.

**Figure 2:**
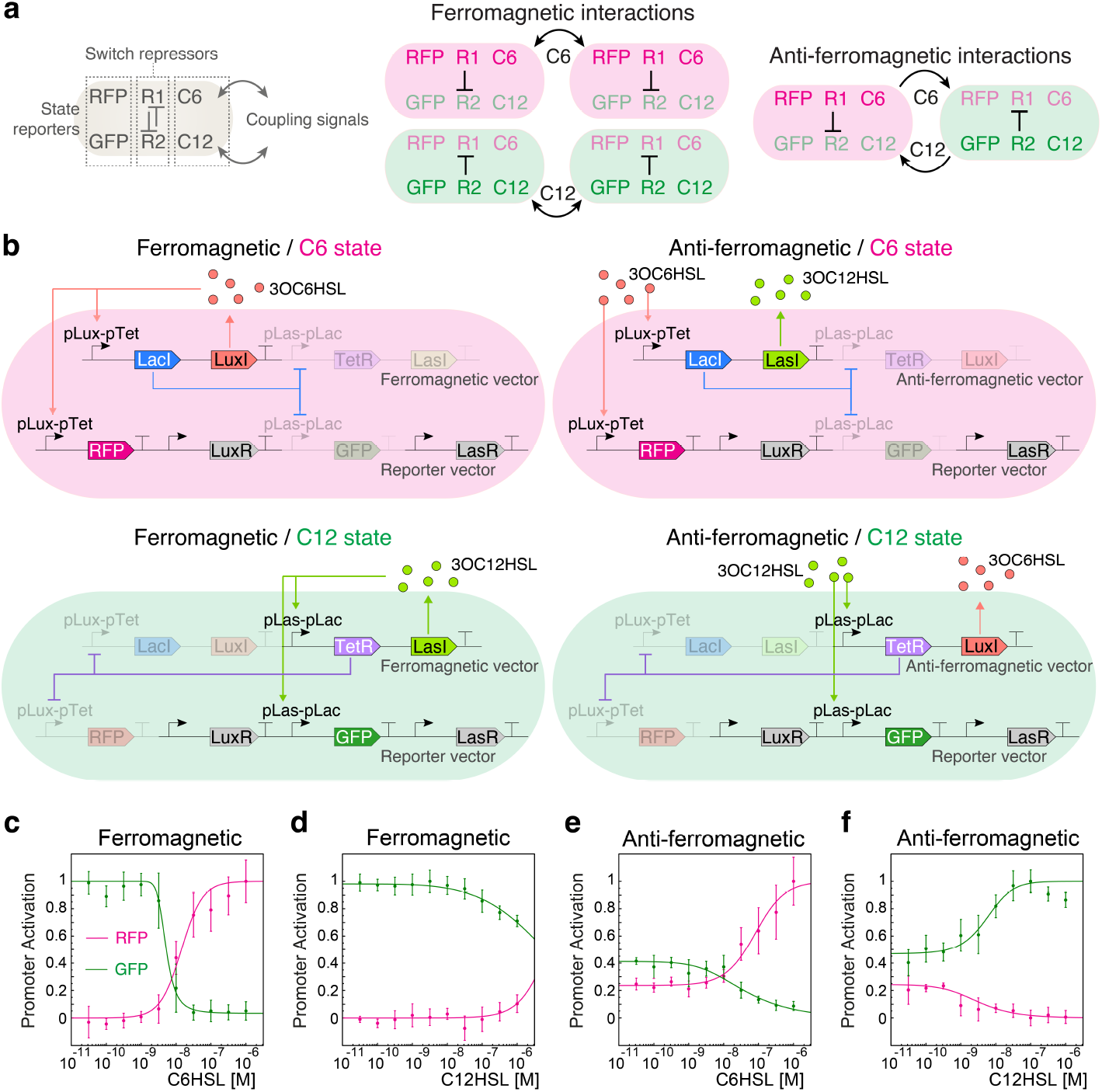
Ferromagnetic and anti-ferromagnetic configurations of coupled, bi-stable synthetic gene networks. (a) Schematic representation of ferromagnetic and anti-ferromagnetic interactions between two neighboring cells. The states, defined by the expression of a red (mCherry2 labeled as “RFP” for simplicity) or green (sfGFP, labeled as “GFP” for simplicity) fluorescent proteins, are determined by mutually inhibiting repressors R1 and R2. Cell states are coupled, in ferromagnetic and anti-ferromagnetic configurations, with neighboring cells by diffusive signals C6 and C12. (b) Gene network arrangement of ferromagnetic and anti-ferromagnetic systems in C6 and C12 states. Ferromagnetic and anti-ferromagnetic systems are composed of a ferromagnetic or anti-ferromagnetic vector and a reporter vector. LasR-C12 and LuxR-C6 complexes were omitted for simplicity. (c-f) Red and green fluorescent protein synthesis rate of *E. coli* cells carrying ferromagnetic (c,d) and anti-ferromagnetic (e, f) systems, grown in liquid medium supplemented with different concentrations of C6HSL (left) and C12HSL (right). Points and error bars correspond to the mean of the fluorescent protein synthesis rates normalized by its maximum value reached in each system and the standard deviation of 4 biological replicates, while lines correspond to the fitting of Eq. 2.

To test the bi-stability condition and coupling properties of the ferromagnetic and anti-ferromagnetic systems, we calculated the red and green fluorescent protein synthesis rate Rudge *et al* (2016) in *E. coli* cells grown in different concentrations of the coupling signals C6HSL and C12HSL (Fig. 2c-f and Additional file 1, Fig. S2). To model SGNs behavior, we considered a simplified version in which each dual promoter (pLux76pTet and pLas81pLac) has two states: an active state that allows the expression of the fluorescent protein gene (ON state), and an inactive state with no expression of this gene (OFF state) Keymer *et al* (2006). The probability *P* of finding each promoter in the active state at equilibrium is:

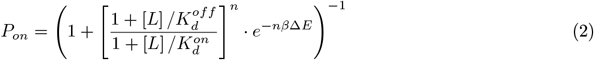

where *L* is the concentrations of the coupling signal (the ligand), Δ*E* = *E*_*off*_ − *E*_*on*_ represents the energy difference between inactive and active promoter without ligand bound, 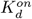 and 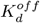are the dissociation constants that characterize the binding of ligands to active and inactive promoters, respectively; and *n* is a Hill exponent to represents cooperative binding Keymer et al (2006).

In a population of cells carrying only the reporter vector, the probability of finding *RFP* and *GFP* promoters active in absence of the inducers C6HSL and C12HSL is close to zero, and it increases as the concentration of its inducer increases (Additional file 1, Fig. S2). This behaviour is reflected in the negative values of Δ*E* and that 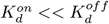 (Additional file 2, Table S2). At very low concentrations of inducers, a population of ferromagnetic cells is only in the green state (Fig. 2c, d). This means that the probability of finding the *GFP* promoter active is 1 (Δ*E >* 0), while the probability of finding the *RFP* promoter active is 0 (Δ*E <* 0) (Additional file 2, Table S3), suggesting that the system is biased towards the production of C12HSL. Accordingly, the pLas promoter has been shown to have a higher basal expression than the pLux promoter Kylilis et al (2018); Grant et al (2016). Since in this system the pLas81pLac promoter directs the expression of *LasI*, its higher basal expression drives cells to produce basal amounts of C12HSL, generating a population of cells in the same state. This also explains why the external addition of C12HSL did not induce a major change in the synthesis rates of fluorescent proteins, except at very high concentrations of this inducer (Fig. 2d) at which signal crosstalk starts to play a relevant role in the activity of the promoters Grant et al (2016). At around 10^−8^ M of C6HSL, there is a drastic decrease in the probability of finding the *GFP* promoter active, which is accompanied by an increase in the probability of finding the *RFP* promoter active (Fig. 2c; compare the values of 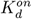 and 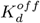 for each promoter in Table S3). Thus, depending on the concentration of C6HSL in the medium, a population of cells carrying the ferromagnetic system can be in one of 3 states: all cells in the green state (low C6HSL), all cells in the red state (high C6HSL), or a mix of red and green cells (around 10^−8^ M of C6HSL). These results suggest that a population of ferromagnetic cells can change between red and green states depending on the concentration of C6HSL in the medium.

In a population of cells carrying the anti-ferromagnetic system, the probability of finding the *GFP* and *RFP* promoters active is different from zero at very low concentrations of inducers (Fig. 2e, f), with values of Δ*E* closer to 0 (Additional file 2, Table S4). In this condition, a population of anti-ferromagnetic cells is in a mixed state, with red and green cells. Microscope analysis of cells in the mixed state revealed that there is no “yellow” cells (data not shown), indicating that individual cells can only be in red or green state at a time. Since *LuxI* is under the control of the pLas81pLac promoter, cells produce a basal amount of C6HSL. Contrary to the case of the ferromagnetic system, this induces the opposite state in other cells, inducing them to produce C12HSL and leading to a balance in the production of both coupling signals. Increasing the concentration of C6HSL in the medium induces an increase in the probability of finding the *RFP* promoter active and a decrease in the probability of finding the *GFP* promoter active (Fig. 2e), while increasing the concentration of C12HSL produces the opposite effect (Fig. 2f). These results show that a population of anti-ferromagnetic cells can change from a mixed state to a population in a red or green state depending on the concentration of the coupling signals in the medium.

These results suggest that both ferromagnetic and anti-ferromagnetic systems are able to couple states in liquid culture, a required property for the emergence of Ising-like pattern when cells are spatially distributed (i.e. grown in solid media) (Fig. 1).

### Ising-like patterns in ferromagnetic and anti-ferromagnetic populations

To test whether the SGNs were able to achieve Ising-like patterns of gene expression such as those observed in CPIM simulations, we studied the fluorescent patterns that emerge in colonies of *E. coli* cells carrying the reporter and ferromagnetic or anti-ferromagnetic vectors. In order to discard any bias related to the properties of the reporters, we constructed another version of the reporter vector in which the promoters directing the expression of the red and green fluorescent proteins are swapped (Additional file 1, Fig. S3). To counteract the higher basal expression of the promoter induced by C12HSL and make red and green states equally likely, ferromagnetic and anti-ferromagnetic, in Figure S4, cells were grown on solid medium supplemented with different concentrations of C6HS. Consistent with what was observed in the data from liquid well plates shown in Figure 2c-f, ferromagnetic colonies grown on solid agar were found to be only in one state in absence (or at very low concentrations) of C6HSL: green for the reporter vector 1 and red for the reporter vector 2 (Additional file 1, Fig. S4A). However, growing ferromagnetic cells on agar plates with 10^−8^ M of C6HSL led to the generation of spatial patterns of red and green cellular state domains across the colonies (Additional file 1, Fig. S4A and Fig. S5), in accordance with the state transition found in liquid well plates (Fig. 2c). A high concentration of C6HSL (10^−8^ M) also produced colonies in only one state but opposite to that of colonies grown at low concentrations of the inducer (Additional file 1, Fig. S4A). At this point, global exogenous concentration of C6HSL appeared to dominate the system over the cell-cell coupling between networks.

Anti-ferromagnetic colonies showed a spatial pattern of red and green domains in the absence (and at very low concentrations) of C6HSL (Additional file 1, Fig. S4B). Under this condition, the center of colonies was dominated by cells in green (reporter vector 1) or red state (reporter vector 2), while the periphery was mainly composed of cells in the opposite state. Due to the higher basal expression of the pLas81pLac promoter, all anti-ferromagnetic cells that give rise to colonies are mostly in the same state. Although these cells synthesize basal amounts of C6HSL, it is not enough to counteract the effects of the basal expression of the promoter in neighboring cells to induce the opposite state, generating a sector of the colony dominated by cells in one state. At some point during the growth of the colony, the C6HSL accumulated in the medium allows cells to change states regardless of promoter basal expression, generating a sector with a mix of red and green domains. As cells continue synthesizing C6HSL, it accumulates to a concentration that triggers a ring of the opposite state in newly born cells in the periphery of the colony. As in ferromagnetic colonies, red and green cellular state domains emerged across the whole colony at 10^−8^ M of C6HSL (Additional file 1, Fig. S4B, and Fig. S5). Compared to those patterns in ferromagnetic colonies at the same C6HSL concentration, the red and green domains generated in anti-ferromagnetic colonies are much smaller. A further increase in the concentration of C6HSL in the medium (10^−8^ M) only produced colonies in the red (reporter vector 1) or green (reporter vector 2) state (Additional file 1, Fig. S4B).

These results show that ferromagnetic and anti-ferromagnetic SGNs allow the self-organization of distinctive patterns in *E. coli* colonies, as partially anticipated by the CPIM simulations (Fig. 1c). However, the formation of fractal-like jagged patterns, characteristic of rod-shaped non-motile *E. coli* cells, caused these patterns to visually differ from those obtained with simulations. These fractal patterns are the result of both mechanisms at play: the chemical coupling and the buckling instabilities generated by the polar cell shape that propagate due to the uni-axial cell growth and division Rudge et al (2013).

In order to analyze the pattern generated by the ferromagnetic and anti-ferromagnetic systems without the influence of uniaxial cell growth, we used the *E. coli* mutant strain KJB24 that forms spherical cells. This strain performs cell division in any direction due to a mutation in *RodA* Begg and Donachie (1998), removing the cell polarity-driven buckling instabilities that give rise to jagged patterns Rudge et al (2013). As observed in colonies of rod-shaped cells, C6HSL increases only the red or green fluorescence of colonies of spherical cells carrying only the reporter vector 1 or 2 (Additional file 1, Fig. S6), respectively. In accordance with the findings on colonies of rod-shaped cells, ferromagnetic Ising-like patterns emerged in colonies of ferromagnetic spherical cells when they were grown in the range of 10^−8^ M of C6HSL, regardless of the reporter vector used (Fig. 3). These patterns look qualitatively more similar to those observed in CPIM simulated ferromagnetic populations than the patterns generated by rod-shaped ferromagnetic cells. Anti-ferromagnetic colonies of spherical cells showed a characteristic pattern of small domains of red and green states at 10^−8^ M of C6HSL (Fig. 3).

**Figure 3:**
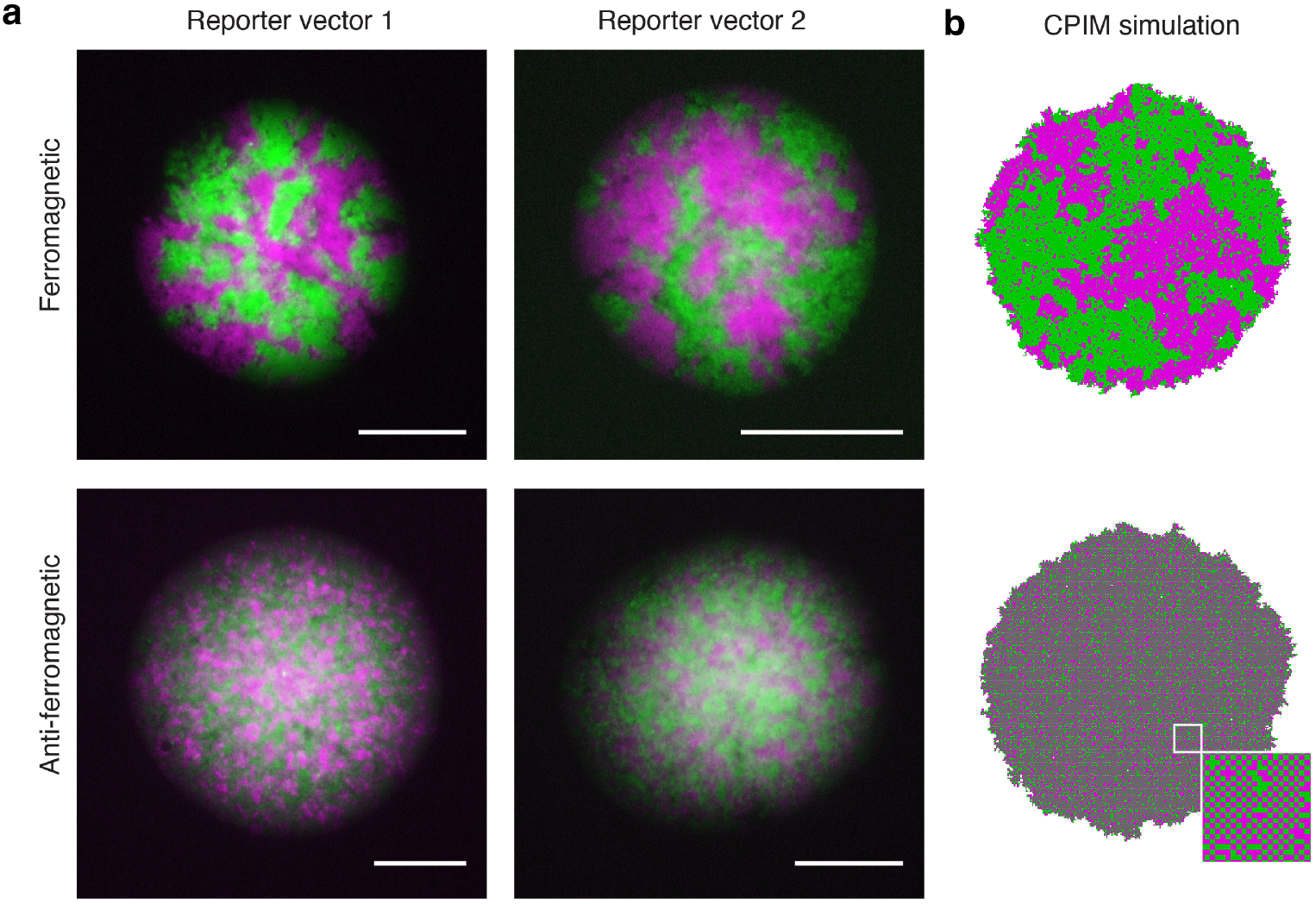
Self-organized patterns of cellular states in ferromagnetic and anti-ferromagnetic colonies. a) Representative images of red and green fluorescent protein patterns that emerge in colonies of spherical *E. coli* cells carrying the ferromagnetic or anti-ferromagnetic systems with reporter vector 1 or 2. Cells were grown on solid M9-glucose medium supplemented with 10^−8^ M of C6HSL, a concentration that counteracts the bias introduced by the basal expression of the pLas81pLac promoter. Images were taken approximately 18 hours after inoculation. Scale bars 100 *µ*m. Red cells are shown in magenta. b) Images obtained from populations simulated with CPIM are included for comparison.

To compare the patterns that emerged in colonies to those observed in simulated populations, we calculated the Hamming distance (Additional file 1, Fig. S7) between colonies of ferromagnetic spherical cells and simulated ferromagnetic populations around the critical value of *T* (*T*_*c*_ = 2.27) (Additional file 1, Fig. S7A, magenta dots). The Hamming distance between two images of equal size is the number of pixel positions at which the values of those pixels are different. Therefore, the smaller the Hamming distance between two images, the more similar those images are. As observed before, a strong coupling between cells (*T < T*_*c*_) leads to the generation of populations with large homogeneous domains, with the same probability of finding populations mostly in red or green state (Additional file 1, Fig. S7B). This explains the great variability observed in the average Hamming distance below the critical value of *T* (Additional file 1, Fig. S7C). Interestingly, the smallest average Hamming distance was found with respect to simulated populations close to the critical value of *T* (Additional file 1, Fig. S7C). These results suggest that the patterns observed in ferromagnetic colonies of spherical cells are similar to simulated populations near the critical transition, at which spatial correlations increase.

To demonstrate that the patterns observed in colonies depend on the coupling signals, we also studied colonies in which red and green states were determined by constitutive expression promoters without chemical coupling between SGNs. These states were located in plasmids that irreversibly segregate to daughter cells after cell division Nunez et al (2017), enabling cells to acquire one of the two states perpetually and creating state domains that only enlarge by cell division. Unlike the patterns observed in ferromagnetic and antiferromagnetic colonies, these colonies exhibited radial domains of segregating sectors such as those observed in Rudge et al (2013) (Additional file 1, Fig. S8). Next, to test whether the observed patterns depend on the internal genetic switch module, we grew ferromagnetic colonies in the presence of IPTG and aTc to inhibit the action of LacI and TetR repressors, respectively. These colonies exhibit only one state, corresponding to the inactive repressor, whereas inhibiting both repressors led cells to adopt the state dictated by the coupling signals they sense (Additional file 1, Fig. S9). These findings suggest that the observed ferromagnetic and antiferromagnetic patterns are the result of coupled and bi-stable gene networks.

### Spatial correlations, cluster size distributions and critical exponents of ferromagnetic and anti-ferromagnetic colonies

To quantitatively compare the patterns generated in ferromagnetic and anti-ferromagnetic colonies of rodshaped and spherical cells, we calculated the spatial autocorrelation function (sACF) Walter (2017) (Fig. 4a and b). The sACF describes how the correlation between two microscopic variables (e.g. the state of each cell in a colony) changes on average as the separation between these variables changes Nounou and Bakshi (2000), allowing the calculation of the characteristic size of the cellular state domains that emerge in a colony.

**Figure 4:**
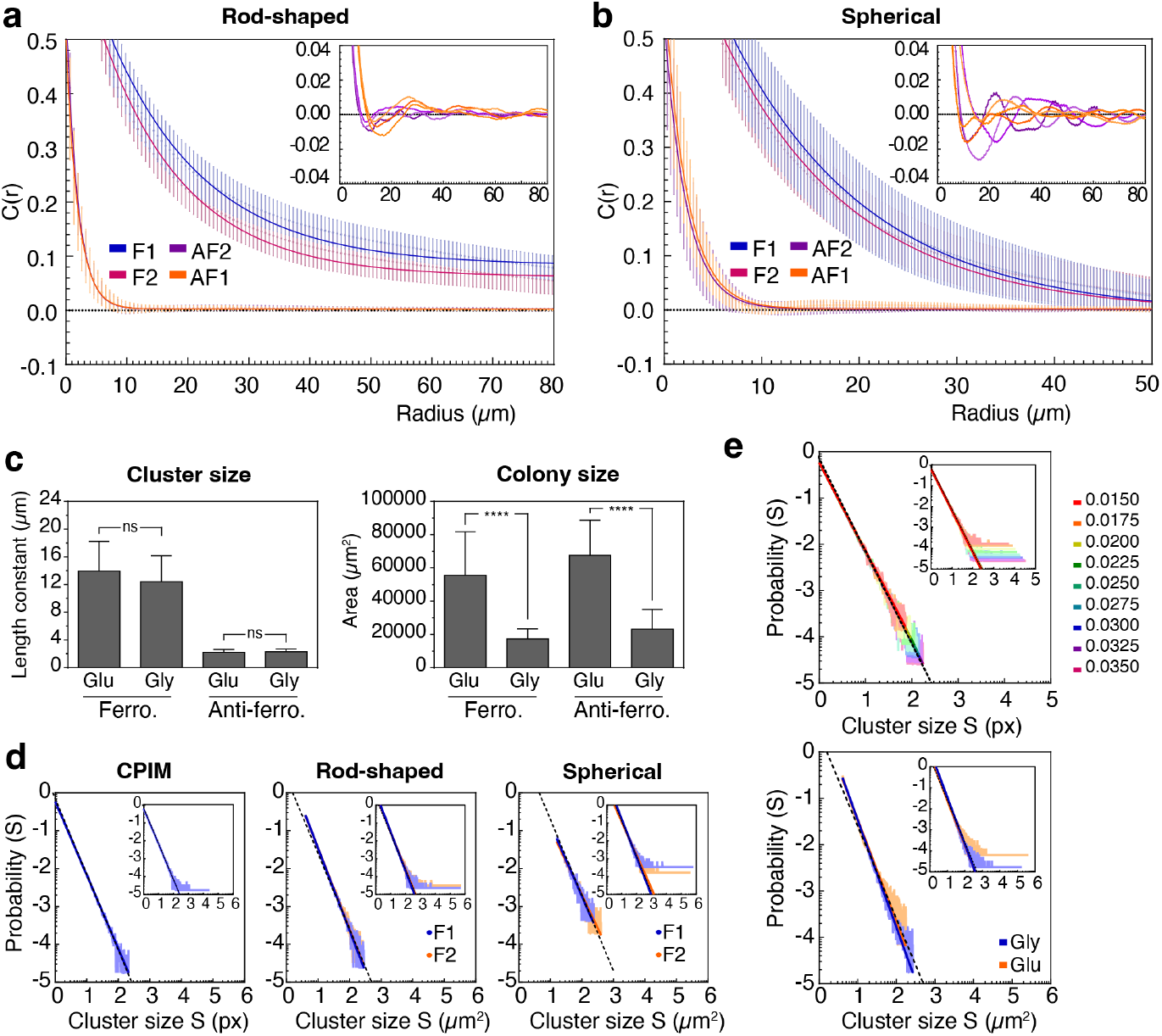
Spatial correlation in ferromagnetic and anti-ferromagnetic colonies. Spatial autocorrelation function *C*(*r*) in colonies of rod-shaped (a) and spherical (b) *E. coli* cells carrying the ferromagnetic (F) and anti-ferromagnetic (AF) systems with reporter vector 1 (F1 and AF1) or 2 (F2 and AF2). Points and error bars correspond to the mean ± the standard deviation of around 40 colonies for each system, and lines correspond to the best fit of the exponential decay equation *y* = *y*_0_ ∗exp(−*x/b*) + *C* to the data. Insets show the oscillating behavior of the sACF around zero of individual anti-ferromagnetic colonies, which is lost when the data is averaged. (c) Length constant and colony size of ferromagnetic and anti-ferromagnetic colonies of spherical *E. coli* cells grown in M9 solid medium supplemented with glucose (Glu) or glycerol (Gly), showing that cell division rate does not affect the spatial correlations. Statistical analysis was performed using an unpaired two-tailed Mann-Whitney test (*α* = 5%). ns (not significant): P *>* 0.05; ∗ ∗ ∗∗ ≤ : P 0.0001. (d) Probability distribution *P* (*S*) (log_10_ − log_10_ plots) of the cluster size *S* (in pixels) for ferromagnetic populations simulated with CPIM (left) and ferromagnetic colonies of rod-shaped (middle) and spherical (right) cells. (e) Probability distribution of the cluster size for ferromagnetic populations simulated with CPIM at different cell birth rates, from 0.0350 to 0.0150 (top), and ferromagnetic colonies of spherical cells grown in glycerol (blue) or glucose (red) (bottom). Plots in (d) and (e) were obtained using the algorithms r_plfit with the default low-frequency cut-off Hanel et al (2017). Solid lines correspond to the power-law 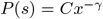 found by the algorithm. Insets show the probability distribution of all the clusters found in the populations (without cut-off), with solid lines corresponding to the best fit of the data to equation *P* (*s*) = *A* ∗ *s*^−*γ*^ found by the least squares method. Dotted lines correspond to a curve with *γ* = 2.00.

In rod-shaped and spherical cells, the correlation function curve decays much faster for anti-ferromagnetic colonies (Fig. 4a, and b), suggesting that the average distance at which two cellular states correlates is shorter in these colonies. Individual colony analysis revealed that most of these colonies show negative values on the correlation curve, indicating the existence of short-range anti-correlations (insets of Fig. 4a, and b). To obtain the characteristic size of the cellular state clusters, we fitted the exponential decay equation *y* = *y*_0_^∗^exp(−*x/b*)+*C* to the data obtained from the computation of the sACF. In this equation, *b* corresponds to the length constant, which is an estimation of the mean size of cellular state clusters that emerge in the colonies. As suggested by the patterns observed in the colonies (Fig. 3), the mean size of the cellular state domains that emerge in ferromagnetic colonies is larger than those observed in anti-ferromagnetic colonies (Table 1). This difference was independent of the reporter vector (R.V.) used. In colonies of rod-shaped cells, the mean size of the clusters in ferromagnetic colonies is approximately 8.1 (R.V.1) and 7.7 (R.V.2) times larger than the mean size in anti-ferromagnetic colonies, while in colonies of spherical cells is approximately 6.7 (R.V.1) and 6.6 (R.V.2) times larger. These results show that the ferromagnetic system leads to larger spatial correlations than those generated by the anti-ferromagnetic system.

**Table 1:**
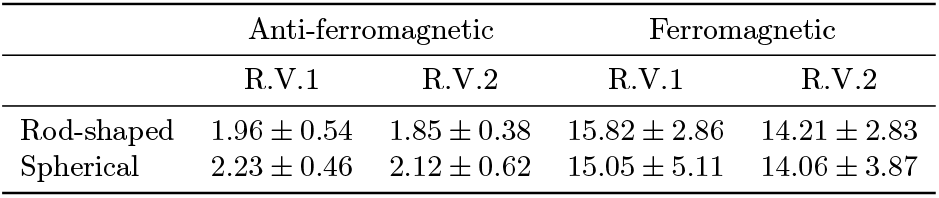
Length constants of ferromagnetic and anti-ferromagnetic colonies of rod-shaped and spherical *E. coli* cells

|  | Anti-ferromagnetic |  | Ferromagnetic |  |
| --- | --- | --- | --- | --- |
|  | R.V.1 | R.V.2 | R.V.1 | R.V.2 |
| Rod-shaped | $1.96 \pm 0.54$ | $1.85 \pm 0.38$ | $15.82 \pm 2.86$ | $14.21 \pm 2.83$ |
| Spherical | $2.23 \pm 0.46$ | $2.12 \pm 0.62$ | $15.05 \pm 5.11$ | $14.06 \pm 3.87$ |

Interestingly, no significant differences were found between the length constants of ferromagnetic colonies of rod-shaped and spherical cells (P value: 0.0557 for F1 rod vs F1 sph and 0.4303 for F2 rod vs F2 sph) (Additional file 1, Fig. S10; Table 1), despite their different qualitative appearance (Fig. 3). This suggests that the spatial correlation that emerges from ferromagnetic SGNs is independent of cell shape and mechanically-driven cell ordering. To further study the robustness of these patterns, we tested whether cell division rate affects the correlation length of Ising-like colonies. CPIM simulations predicted that, while the size of the populations decreases with the value of the birth rate, no significant differences were found in the length constants when the birth rate decreases from 0.0255 to 0.0180 (P value: 0.8895 for 0.0250 vs 0.0255, 0.4107 for 0.0225 vs 0.0255, 0.2890 for 0.0200 vs 0.0255, 0.8436 for 0.0195 vs 0.0255, 0.1402 for 0.0190 vs 0.0255, 0.0502 for 0.0185 vs 0.0255, 0.0606 for 0.0180 vs 0.0255). Between these values of birth rate, the size of the population is reduced to approximately 0.48 (Additional file 1, Fig. S11). To test this model prediction, we analyzed spatial correlations in colonies of spherical ferromagnetic and anti-ferromagnetic cells grown in minimal solid medium supplemented with glucose or glycerol as the carbon source. It has been observed that the generation time of *E. coli* cells increase when grown in a glycerol-supplemented medium compared to cells grown in a glucose-supplemented medium Taheri-Araghi et al (2015). In agreement with the CPIM predictions, length constants were no significantly affected in ferromagnetic and anti-ferromagnetic colonies that had reduced their size by more than half (P value cluster size: 0.1453 for Glu vs Gly Ferro, 0.7395 for Glu vs Gly Anti-ferro; P value colony size: *<* 0.0001 for Glu vs Gly Ferro, *<* 0.0001 for Glu vs Gly Anti-ferro) (Fig. 4c). Together, these results suggest that the scaling properties of Ising-like, bi-stable, and coupled SGNs are independent of the cell shape and division rate.

To further characterize the behavior of the ferromagnetic system, we analyzed the size distribution of the cellular state clusters. As seen in Figure 4d, the probability distribution *P* (*S*) of cluster size *S* for ferromagnetic colonies of rod-shaped and spherical cells shows a scale invariant distribution of the form *P* (*s*) ∼*s*^−*γ*^. The values of *γ* calculated for the ferromagnetic systems are consistent with the exponent of cluster size distribution near the critical percolation threshold, which follows a power-law decay with an exponent of 2.055 Stauffer and Aharony (1994) (Rod-shaped: R.V.1 = 2.17, R.V.2 = 2.14. Spherical: R.V.1 = 1.91, R.V.2 = 1.76) (Additional file 2, Table S5). This exponent was also consistent with the value of *γ* for simulated populations with ferromagnetic interactions at *T* = *T*_*c*_ (*γ* = 1.94). Interestingly, the simulations also showed that the size distribution of cellular state clusters is not affected by changes in the birth rate (Fig. 4e top) (Additional file 2, Table S6). As predicted by CPIM simulations at different cell growth rates, similar power law distributions were found for ferromagnetic colonies grown in glycerol or glucose, with scaling exponent *γ* equal to 2.19 for glucose and 2.31 for glycerol(Fig. 4e bottom) (Additional file 2, Table S5). This suggests that the scaling properties are maintained upon changes in cell growth rate within the evaluated range.

## 3 Discussion

Understanding how gene spatial correlations emerge from interacting genetic networks is a fundamental problem in biology. Guided by a two-dimensional (2-D) Contact Process (CP) model incorporating Ising mechanisms to represent gene expression, we show how Synthetic Gene Networks (SGNs) with two states, which are positively or negatively coupled, give rise to a rich repertoire of shortand long-range correlations (and anticorrelations). These SGNs are capable of self-organizing into long-range correlations with power-law scaling properties or checkerboard-like patterns similar to ferromagnetic and anti-ferromagnetic configurations of the Ising Model (IM) near critical points, respectively. Near the critical point, the spatial autocorrelation function of simulated “ferromagnetic populations” follows a power-law decay with an exponent consistent with the value of the IM at the critical temperature. On the other hand, the scaling exponent γ calculated for both simulated and *in vivo* ferromagnetic colonies were close to the exponent of the cluster size distribution near the critical percolation threshold (Larkin et al (2018); Stauffer and Aharony (1994)). At this critical point, the system moves from a regime of only localized short-range patches to one with clusters that span the entire system.

To further understand the behavior at the critical point as to investigate the value of the critical exponents of the CPIM model, we performed a finite-size scaling analysis (see Additional file 3, Supplementary Note 2: Finite-size scaling analysis; Additional file 2, Fig. S23; Additional file 2, Table S7) for the demographic (birth/colonization or extinction/death) parameters used to model our experimental colonies (*b* = 0.03 and *d* = 0.00001). This corresponds to a CP of high reproductive number (*R*_0_ = *b/d* ≫ 1, Additional file 2, Fig. S23A) where most lattice sites are occupied by spin states (−1 or +1). Our analysis suggests that the CPIM is not in the same universality class as the Ising model (IM), although both the height of the peak of magnetic susceptibility per occupied site *X*^*N*^ and the average magnetization per occupied site 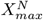 at the critical point 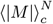 scale with the size of the lattice in a similar way the IM scales with system size (exponents *γ* and *β*) but differently with respect to the scaling to system size of Weber and Buceta’s model (WB; Weber and Buceta (2016)) as shown in Table S7 (Additional file 2). Moreover, the critical temperature in the thermodynamic limit *T*_*c*_ found for the CPIM is roughly the same as the one found for the 2-D IM Ibarra-García-Padilla et al (2016). Note however that both the exponents and the critical temperature might be different for different CP parameters. As shown in Figure S19B, if we increase the death (or extinction) rate *d*, the CPIM shifts its *T*_*c*_ to lower values. Further theoretical analysis of the CPIM could investigate such phenomena elsewhere as it is beyond the scope of our work, which is mostly focusing on understanding our experimental colonies. Follow-up experiments modulating demographic parameters could be achieved using microfluidic devices in order to test these predictions of the CPIM model. In this study, however, we have focused on quasi-2D colonies growing on an agar surface as an introduction to our SGNs. An interesting result of our finite-size scaling analysis is the fact that the position of the magnetic susceptibility per occupied site peak 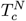 does not scale with the system size in the same way that the IM scales (Additional file 2, Table S7), exponent *ν*). This deviation from the IM behavior is consistent with the observation that the temperature at which the magnetic phase transition occurs in our model does indeed depend on colonization/extinction dynamics (i.e. the CP). For increased values of *d*, extra vacancy introduces noise in the IM behavior, and thus stronger coupling or lower temperature are needed to compensate.

These theoretical results combined with our experimental ones suggest that the “ferromagnetic” genetic system can pose colonies near the critical point of a phase transition in which far regions in the colony are correlated. These findings are in agreement with the theoretical work of WB Weber and Buceta (2016), who found that simulations of toggle switches with coupled states can exhibit phase transitions described by the theory of critical phenomena (Weber and Buceta (2016)) (see Supplementary Note 4, Additional file 4). Although our results suggest that the behavior of these SGNs could belong to a universality class (IbarraGarcía-Padilla et al (2016); Goldenfeld (1992)) related to the one of the IM, the lack of a control parameter in the *in vivo* experimental system limits the search for universal critical exponents in the experimental colonies. In our lattice model, the control parameter T (in comparison to the value of J) accounts for how strongly coupled sites are in terms of responding to the state of their neighbors. Such a control parameter could be realized *in vivo* by the regulation of coupling signal levels outside the cells or their transport rate across the cell membrane (Weber and Buceta (2016)). To evaluate how a higher transport of autoinducers can affect our results, we also investigated a CPIM where the Ising coupling interactions take place in a Next Nearest Neighborhood (NNN). As predicted by the mean-field theory of the IM, if the number of interacting neighbors increases, the critical temperature *T*_*c*_ shifts to higher values (Additional file 2, Fig. S19A).

Another interesting observation is that although the CPIM considers flipping rates between spin states which only depend on local energy differences, the dynamics of the empirical pattern of gene expression of our SGNs within the growing colonies varies with both spatial coordinates within the colony as well as its age. When comparing colonies at 14, 18, and 22 hours post inoculation (see Additional file 3, Supplementary Note 3: Time series analysis; and Additional file 2, Figure S18) we see that as cells within the middle of the colony approach stationary phase conditions patterns slow down their dynamics.

Although in our theoretical approach we modeled the behavior of gene expression as a function of inducer concentration (see Fig. 2c-f; Additional file 1, Fig. S4, and Eq.2) we did not extend this model to the CPIM. The reasoning behind this decision is to avoid assumptions about the microscopic dynamics of transport, conceiving a very generic model combining the CP to model our demographic condition (a quasi-2D growing colony) with the actual IM as a model of a coupled toggle switch. The approach of WB Weber and Buceta (2016) offers a heuristic complement to study the microscopic model. We trust that our empirical system will encourage further experimental work in microfluidics devices where the transport of coupling molecules, as well as demographic parameters, can be controlled. In such a controlled environment, where cells are kept in log phase, further theoretical work can be developed. The most striking result hinting that our model captures the essential dynamics of our experiments with SGNs is the fact that it reproduces empirical patterns in both, ferromagnetic as well as antiferromagnetic colonies (see Fig. 3). The main purpose of our modeling efforts is to understand the induction of our genetic constructs (Equation 2) and to interpret our experiments with SGNs in growing colonies (CPIM).

The generation of colonies with all SGNs in the same state above a critical concentration of the C6HSL coupling signal indicates that these SGNs can also align their dynamics to a global exogenous force. This exogenous force can be interpreted as an analog to the externally applied magnetic field, represented by the second term in the Hamiltonian of the Ising model Ibarra-García-Padilla et al (2016); Selinger (2016); Noble et al (2018). While the interaction energy favors the alignment between spins, the field energy favors the alignment of all spins with the external field. A high concentration of the coupling signal counterbalances the effect of local coupling interactions and favors the alignment of SGN state with this external C6HSL field. The interplay between these two mechanisms has also been observed in spatial ecology, where fractal longrange correlations in the yield of pistachio trees, which emerge due to coupling interactions between trees, becomes homogeneous under global exogenous forces such as weather Noble et al (2018), a phenomenon the authors related to the widely known Moran effect Moran (1953). The incorporation of a magnetic field effect to CPIM would allow to integrate the contribution of endogenous coupling interactions and global exogenous forces that tend to synchronize the entire population. To further explore this avenue of research, future work will also seek to implement an *in vivo* field, whose level and regulation will be orthogonal to the mechanisms controlling bistability and coupling in the current designs (i.e. HSLs).

This work shows how two-state switches with different coupling mechanisms can lead to spatial patterns of rich scaling properties in isogenic bacterial populations. Microbes are constantly adjusting their metabolism to changing environments, and even clonal populations can become spatially structured, giving rise to interacting metabolic subpopulations shaped by cellular uptake and release of coupling compounds (Dal Co et al (2019); Rosenthal et al (2018); Cole et al (2015)). Whether local metabolic couplings (e.g. Shapiro (1998); Blanchard and Lu (2015); Mee et al (2014); Cole et al (2015); Rosenthal et al (2018); Liu et al (2015); Dal Co et al (2020)) can lead to Ising-like patterns in natural multicellular systems remains to be explored. This work provides a minimal system to address these questions as well as other fundamental problems in developmental and microbiology, such as phase transition and symmetry breaking (Weber and Buceta (2016)), with promising applications for the engineering of pattern formation, synthetic bacterial consortia, and artificial morphogenesis (Nguyen *et al* (2018); Teague *et al* (2016); Johnson *et al* (2017); Santos-Moreno and Schaerli (2019); Ebrahimkhani and Levin (2021); Davies and Glykofrydis (2020); Bittihn et al (2018); Solé et al (2018)).

## 4 Methods

### 4.1 Computational modeling

The code for the simulation of the Ising model during the growth of a bacterial colony was written in the C programming language, and it is available in Github (https://github.com/jekeymer/Contact-Process-Ising-Model). Additional information on the simulation of the Ising model during quasi-2D colony growth can be found in the Supplementary Note 1, Additional file 3. The graphical user interface was created with GTK+ 3 (https://developer.gnome.org/gtk3/stable/).

### 4.2 Plasmids

All the vectors used in this work, listed in Additional file 2, Table S1, were constructed by Golden Gate Engler et al (2008) and Gibson Assembly Gibson et al (2009). Level 0 modules used in the Golden Gate assembly containing one of the four genetic elements that are part of a Transcriptional Unit (promoter, Ribosome Binding Site RBS, coding sequence, and terminator), were either obtained from the CIDAR MoClo kit Iverson et al (2016) deposited in Addgene (Kit #1000000059) or constructed by Gibson Assembly using gBlocks supplied by IDT (idtdna.com). The pLux76pTet and pLas81pLac double promoters designed in this work were synthesized based on the sequences of pLux76 and pLas81 promoters used in Grant *et al* (2016). The sequence of *mCherry2* was obtained from Shen *et al* (2017). Different Level 1 vectors containing Transcriptional Units were generated by Golden Gate combining different Level 0 modules. Four of these Level 1 Transcriptional Units were combined together by Gibson Assembly to generate the Level 2 reporter and Ising vectors. PCR fragments used in the Gibson Assembly were amplified using Phusion HighFidelity DNA Polymerase (NEB) and were visualized on a blue LED transilluminator (https://iorodeo.com) using SYBR Safe (Thermofisher). The purification of the vectors was performed using the Wizard Plus SV Minipreps DNA Purification System (Promega), while the purification of the PCR fragments was performed using the Wizard SV Gel and PCR Clean-Up System (Promega).

### 4.3 Bacterial strains and Growth conditions

All experiments were performed using the *E. coli* TOP10 (Invitrogen) or KJB24 strains. KJB24 strain contains a stop codon mutation in the cell wall protein RodA, which results in the generation of spherical cells, and a second mutation that allows cells to grow in rich medium Begg and Donachie (1998). To transform cells with Ferromagnetic and Antiferromagnetic systems, cells of TOP10 and KJB24 strains were made competent by the CCMB80 method (http://openwetware.org/wiki/TOP10_chemically_competent_cells). Cells were grown on LB liquid medium (tryptone 10 g, yeast extract 5 g, NaCl 5 g, and distilled water to a final volume of 1 L) or on M9-glucose liquid medium (1x M9 salts supplemented with MgSO_4_∗7H_2_O 2 mM, CaCl_2_ 0.1 mM, glucose 0.4% and casamino acids 0.2%, where 1 L of 5× M9 salts contains 64 g of Na_2_HPO_4_∗ 7H_2_O, 15 g of KH_2_PO_4_, 2.5 g of NaCl, and 5 g of NH_4_Cl), where 1.5% w/v agar was added to the corresponding liquid medium to prepare LB agar or M9-glucose agar medium. When necessary, the medium was supplemented with 50 *µ*g/mL kanamycin, 100 *µ*g/mL carbenicillin, 50 *µ*g/mL spectinomycin or 10 *µ*g/mL chloramphenicol. In order to prepare the stock solutions of acyl-homoserine lactone molecules, 3-oxohexanoyl-homoserine lactone (C6HSL, Cayman Chemicals) and 3-oxododecanoyl-homoserine lactone (C12HSL, Sigma) were dissolved in DMSO to a concentration of 0.067 M. Before being used, both acylhomoserine lactones were first diluted in ethanol to a concentration of 2 mM, and then diluted in M9-glucose medium to the described concentrations. To obtain images of colonies, cells were grown overnight at 37 ^°^C in liquid M9-glucose medium and diluted 1:100 in the same fresh liquid medium. Cells were then grown to an optical density at 600 nm of 0.2, diluted 1:1000 and 20 *µ*L of these dilutions were plated onto M9-glucose agar plates with the appropriate antibiotic and the described concentrations of C6HSL. To compare the effects of growth rate in the emergence of patterns, cells were also plated onto M9-glycerol agar plates, in which the glucose was replaced with 0.2% glycerol.

### 4.4 Plate fluorometry

Cells were grown overnight in a shaking incubator at 37 ^°^C in 4 ml of liquid M9-glucose medium supplemented with the appropriate selective antibiotics. The overnight cultures were diluted 1:1000 in the same fresh liquid medium, and 200 *µ*l of these dilutions were transferred into a well of a 96-well clear-bottom plate and supplemented with the described concentration of C6HSL or C12HSL. The plates were placed in a Synergy HTX plate reader (BioTek) and fluorescence (sfGFP: 485/20 nm excitation, 516/20 nm emission; mCherry2: 585/10 excitation, 620/15 nm emission) and optical density (600 nm) were measured every 10 min for 24 hours. The plates were maintained at 37 ^°^C during the experiment and were shaken at 200 rpm between readings.

### 4.5 Microscopy and Image analysis

A Nikon Ti microscope equipped with 10x, 20x, and 40x objectives, and FITC and TRITC Filter Cube Sets were used to obtain the images of the colonies. Images were acquired using the Nikon NIS-Elements BR software. The processing and analysis of the images was performed using the Fiji distribution of ImageJ Schindelin et al (2012). Single-channel images of the colonies were created by merging a z-stack using the Extended Depth Field plugging, while multi-channel images were merged using the Merge command. Before the analysis of the images, single-channel images were converted into 8-bits, the background was removed using the Subtract Background command and the images were binarized using the Automatic Threshold plugging. To investigate the existence of a characteristic spatial distribution of the cellular-state domains, we used the AutoCorrelation Function (ACF) plugging Walter (2017) (https://github.com/vivien-walter/autocorrelation) to calculate the spatial Autocorrelation Function (sACF) of binarized images of whole colonies. A value of 1 or −1 of the sACF means a perfect correlation or anticorrelation, respectively, while a value of 0 means no correlation. To calculate the mean size of the cellular state clusters generated in the colonies we used Gnuplot (http://www.gnuplot.info/) to fit the data of the sACF to the one phase exponential decay equation *y* = *y*_0_ ∗exp(−*x/b*) + *C*, where *b* is the length constant, which correspond to the average size of the cellular states domains generated in the colonies.

To obtain the probability distribution of cluster sizes, we calculate the number and size of the clusters of binarized images of colonies and simulations using the Find Connected Regions Plugin of ImageJ (http://homepages.inf.ed.ac.uk/s9808248/imagej/find-connected-regions/). The values of the power-law exponents *γ* were estimated by finding the best fit for all the data using the least squares method and also using the matlab implementation of the algorithm r_plfit(k,’hist’) developed by Hanel et al. with default low frequency cut-offs Hanel et al (2017). For the colonies, only clusters greater than 1 *µ*m were considered for the analysis.

To determine the similarity between ferromagnetic colonies and ferromagnetic populations obtained from CPIM simulations, we calculate the Hamming distance using a custom Python program that determines the number of pixel positions in which the images are different. Binarized images of both colonies and simulations were scaled to have the same number of pixels, saved as a binary array in a text file, and the Python program was used to compare each position in the array (which represents a pixel of the binarized image) of two images. If in that position both images have the same pixel value, the program adds a 0, but if the values are different, the program adds a 1. The total number of pixels in which the images are different is then divided by the total number of pixels to obtain the Hamming distance. For this analysis, 42 colonies of ferromagnetic cells with reporter vector 1 and 65 colonies of ferromagnetic cells with reporter vector 2 were compared with 10 simulated populations for each value of the control parameter between 2 and 2.54. To find the smallest value of the Hamming distance between a colony and a simulated population, the image of the simulated population was rotated every 15 degrees, generating a total of 24 versions for each simulated population. Thus, the final value of the Hamming distance corresponds to the smallest value obtained from the calculation of the distance between the colony and each version of the simulated population. One-way ANOVA followed by Dunnett’s multiple comparisons test, and non-parametric, unpaired two-tailed MannWhitney test (*α* = 5%) were performed using GraphPad Prism for Windows, GraphPad Software, San Diego, California USA, http://www.graphpad.com.

All the raw data is available in an open data repository in Zenodo (https://doi.org/10.5281/zenodo. 8121516).

## Acknowledgments

KS was supported by Beca de Doctorado Nacional CONICYT 2016 (21160554); JK was supported by ANID – Núcleo Milenio Física Materia Activa Iniciativa Científica Milenio and ANID Fondecyt Regular 1191893; FF was supported by ANID – Millennium Science Initiative Program – ICN17_022 and ANID Fondecyt Regular 1211218. AL was supported by the VRI and the Ph.D. program in Biological and Medical Engineering from Pontificia Universidad Católica de Chile. Powered@NLHPC: This research was partially supported by the supercomputing infrastructure of the NLHPC (ECM-02). We thank Tim Rudge, Janneke Noorlag, Peter Galajda, Miles Wetherington, Anton Kan and members of FF group for valuable comments and feedback. We thank the two anonymous reviewers for the comments that have helped strengthen the manuscript.

## Author Contributions

FF, JEK, and KS conceived and designed the study. AL and KS performed experiments and data analysis. JEK, KS and AL built the model. FF, JEK, and KS drafted the manuscript. All authors read, edited, and approved the final manuscript.

### List of abbreviations

CPIM: Contact Process Ising Model
SGN: Synthetic Gene Network
C6HSL: 3-oxo-C6-homoserine lactone
C12HSL: 3-oxo-C12-homoserine lactone
RFP: red fluorescent protein
GFP: green fluorescent protein
IPTG: Isopropyl *β*-D-1-thiogalactopyranoside
sACF: spatial autocorrelation function
Gluglucose Gly: glycerol
F: Ferromagnetic
AF: Anti-Ferromagnetic
atc: Anhydrotetracycline
WB: Weber-Buceta model

## 5 Supplementary information

### Additional file 1

**Supplementary figures S1:** Power-law decay of the sACF in ferromagnetic populations simulated using CPIM. **Supplementary figures S2:** Red and green fluorescent protein synthesis rate of *E. coli* cells carrying the reporter vector. **Supplementary figures S3:** Induction of red and green fluorescence by C6HSL in colonies of rod-shaped cells carrying different versions of the reporter vector. **Supplementary figures S4:** Red and green state patterns generated in ferromagnetic and anti-ferromagnetic colonies at different concentration of C6HSL. **Supplementary figures S5:** Cellular state patterns in ferromagnetic and antiferromagnetic colonies. **Supplementary figures S6:** Induction of red and green fluorescence by C6HSL in colonies of spherical cells carrying different versions of the reporter vector. **Supplementary figures S7:** Hamming distance between ferromagnetic colonies of spherical cells and simulated populations with CPIM. **Supplementary figures S8:** Sharp boundaries generated between cellular state domains in colonies of spherical cells due to the segregation of fixed-state reporter vectors. **Supplementary figures S9:** Cellular state switch in ferromagnetic colonies grown on solid media supplemented with inhibitors. **Supplementary figures S10:** Characteristic size of cellular states domains of ferromagnetic and anti-ferromagnetic colonies. **Supplementary figures S11:** Characteristic size of cellular state domains and average size of simulated populations with ferromagnetic interactions.

### Additional file 2

**Supplementary figures S12:** Colonies of rod-shaped *E. coli* cells carrying the ferromagnetic system with reporter vector 1 or 2. **Supplementary figures S13:** Colonies of rod-shaped *E. coli* cells carrying the anti-ferromagnetic system with reporter vector 1 or 2. **Supplementary figures S14:** Colonies of spherical *E. coli* cells carrying the ferromagnetic system with reporter vector 1 or 2. **Supplementary figures S15:** Colonies of spherical *E. coli* cells carrying the anti-ferromagnetic system with reporter vector 1 or 2. **Supplementary figures S16:** Structure of the Model. **Supplementary figures S17:** Neighborhoods and Probability **Supplementary figures S18:** Colony size expansion in a 3 point time series. **Supplementary figures S19:** Magnetization and Neighborhood size. **Supplementary figures S20:** Average size of spherical *E. coli* cells. **Supplementary figures S21:** Time series of colonies of spherical *E. coli* cells carrying the ferromagnetic system with reporter vector 1. **Supplementary figures S22:** Dynamics of reporter gene expression (spin flipping rates) in ferromagnetic colonies of spherical *E. coli* cells. **Supplementary figures S23:** Finite-size scaling analysis of Contact Process Ising Model (CPIM). **Supplementary table S1:** Plasmids used in this study. **Supplementary table S2:** Biological parameters of reporter vector. **Supplementary table S3:** Biological parameters of Ferromagnetic vector **Supplementary table S4:** Biological parameters of Anti-Ferromagnetic vector **Supplementary table S5:** Scaling exponent *γ* of populations with ferromagnetic interactions. **Supplementary table S6:** Scaling exponent *γ* of simulated populations with ferromagnetic interactions at different birth rates. **Supplementary table S7:** Numerical critical exponents and critical temperature of the two-dimensional Ising model, Weber-Buceta model (WB) Weber and Buceta (2016), and the Contact Process Ising Model (CPIM).

### Additional file 3

**Supplementary Note 1:** Simulation of the Ising model during the growth of a quasi-2D colony. **Supplementary Note 2:** Finite-size scaling analysis **Supplementary Note 3:** Time series analysis **Supplementary Note 4:** Comparison of our work with the work of Weber and Buceta

## 6 Ethics approval and consent to participate

Does not apply.

## 7 Consent for publication

All authors consent the publication of this manuscript in BMC Biology.

## 8 Competing interests

The authors declare that they have no competing interests.

## 9 Availability of data and materials

All the raw data is available in an open data repository in Zenodo (https://doi.org/10.5281/zenodo. 8121516). The CPIM code is available in Github (https://github.com/jekeymer/Contact-Process-Ising-Model)

## Notes

### Competing Interest Statement

The authors have declared no competing interest.

### Summary of Updates

Author updated

