## Additional file 1 for "Spatial biology of Ising-like synthetic genetic networks"

July 12, 2023

1. ANID – Millennium Science Initiative Program, Millennium Institute for Integrative Biology (iBio), Santiago, Chile.
2. Institute for Biological and Medical Engineering, Schools of Engineering, Medicine and Biological Sciences, Pontificia Universidad Católica de Chile, Santiago, Chile.
3. Institute of Advanced Studies, Shenzhen X-Institute, Shenzhen, China.
4. Schools of Physics and Biology, Pontificia Universidad Católica de Chile, Santiago, Chile.
5. Department of Natural Sciences and Technology, Universidad de Aysén, Coyhaique, Chile.
6. FONDAP Center for Genome Regulation - Department of Molecular Genetics and Microbiology, Pontificia Universidad Católica de Chile, Santiago, Chile.

Keywords: Ising model, bi-stable, synthetic gene networks, spatial correlation, criticality

### Additional file 2

### Supplementary figures S12-S23 and tables S1-S7

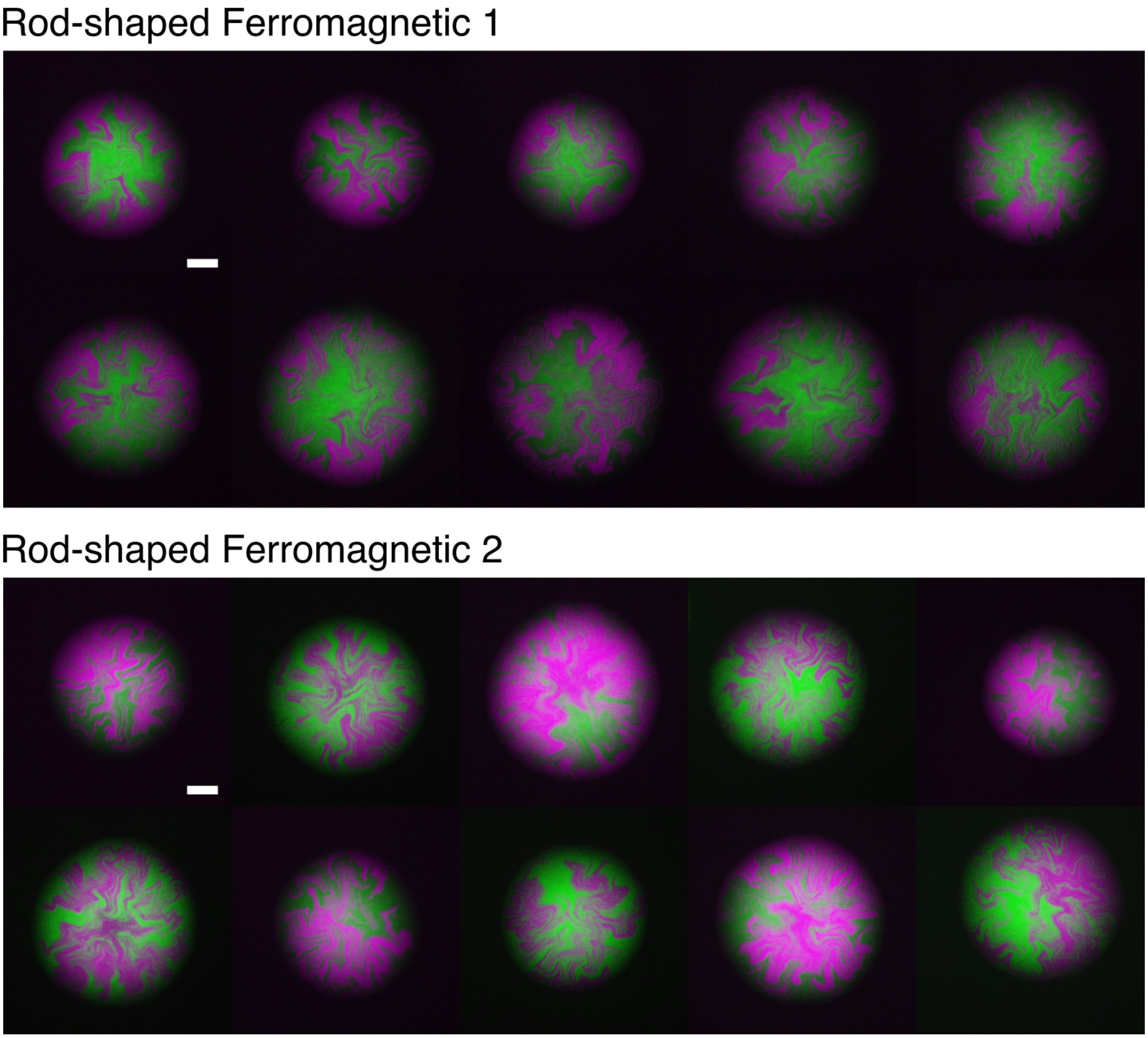

Figure S12: **Colonies of rod-shaped *E. coli* cells carrying the ferromagnetic system with reporter vector 1 or 2.** Cells were grown on solid M9-glucose medium supplemented with $10^{-8}$ M of C6HSL. Images were taken approximately 14 hours after inoculation. Scale bars 100 $\mu$m.

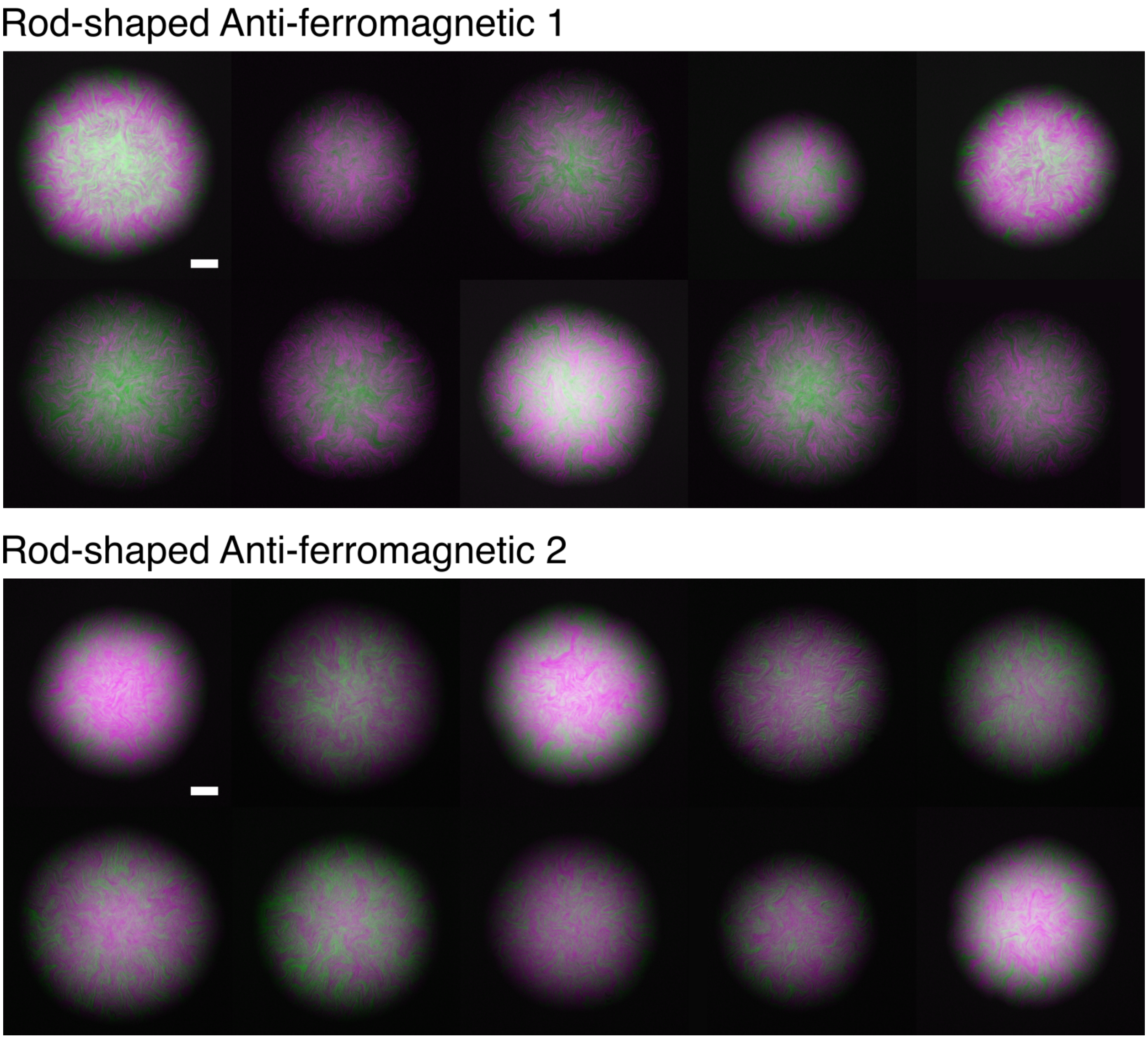

Figure S13: **Colonies of rod-shaped *E. coli* cells carrying the anti-ferromagnetic system with reporter vector 1 or 2.** Cells were grown on solid M9-glucose medium supplemented with $10^{-8}$ M of C6HSL. Images were taken approximately 14 hours after inoculation. Scale bars 100 $\mu$m.

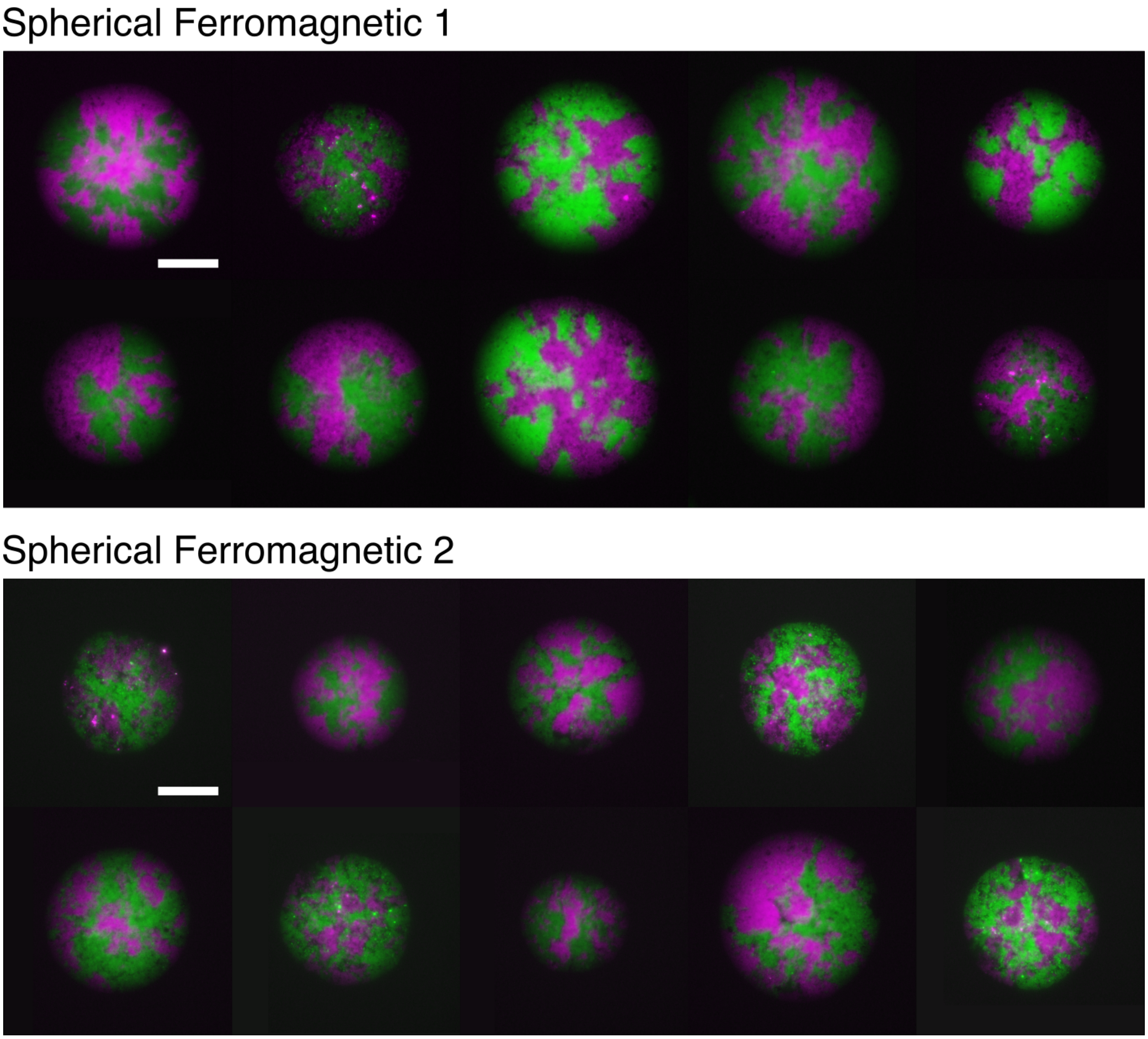

Figure S14: **Colonies of spherical *E. coli* cells carrying the ferromagnetic system with reporter vector 1 or 2.** Cells were grown on solid M9-glucose medium supplemented with $10^{-8}$ M of C6HSL. Images were taken approximately 18 hours after inoculation. Scale bars 100 $\mu$m.

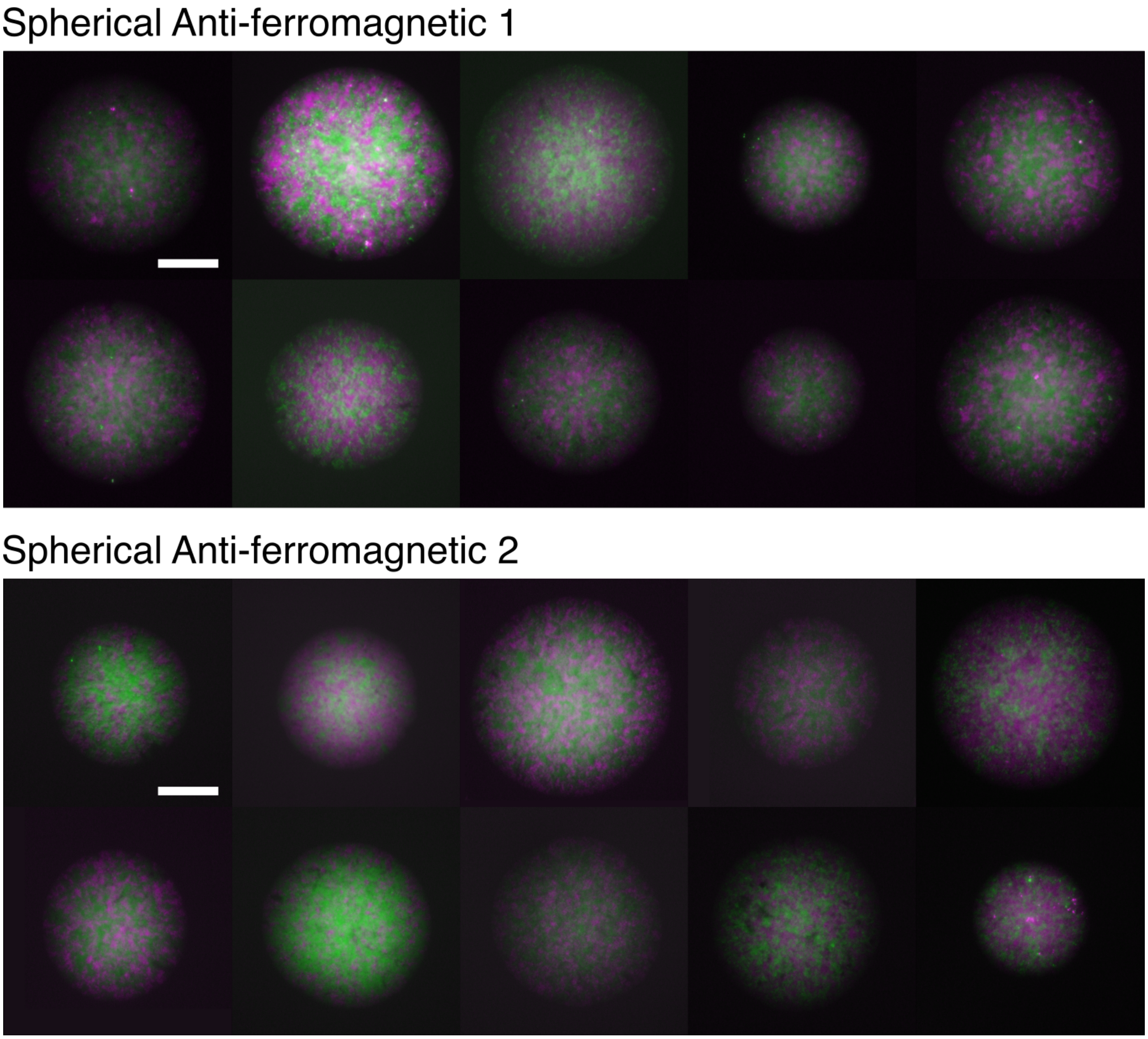

Figure S15: **Colonies of spherical *E. coli* cells carrying the anti-ferromagnetic system with reporter vector 1 or 2.** Cells were grown on solid M9-glucose medium supplemented with $10^{-8}$ M of C6HSL. Images were taken approximately 18 hours after inoculation. Scale bars 100 $\mu$m.

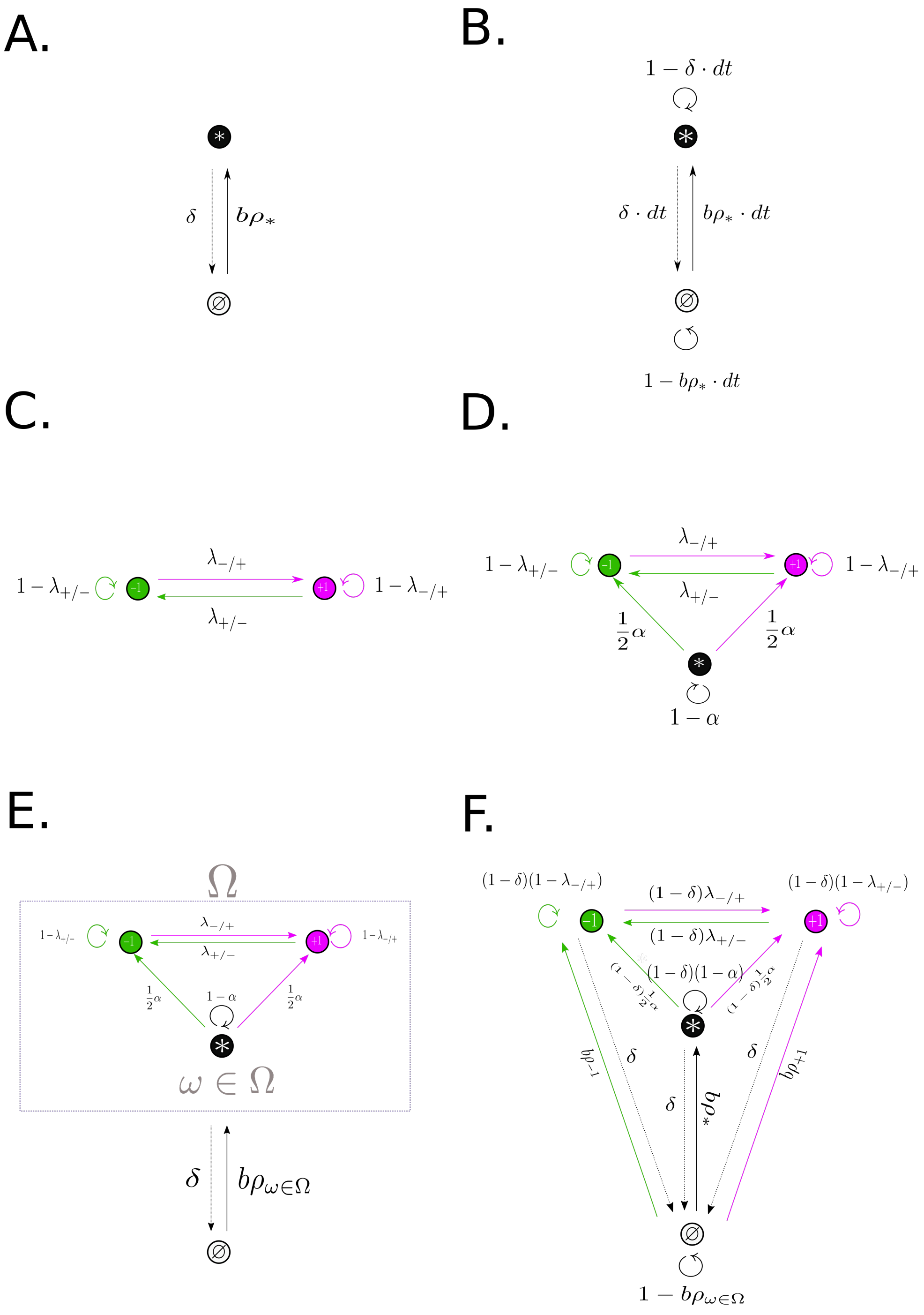

Figure S16: **Structure of the Model.** A) The contact process with birth/colonization rate $b$ and death/extinction $\delta$. B) The contact process as a Markov chain of transition probabilities. C) Ising model steps as a Markov chain, here $\lambda_{-/+}$ and $\lambda_{+/-}$ represent spin-flip probabilities. D) Transition probabilities for an Ising with differentiation process. E) Contact process which includes 3 internal labels of occupancy and where $dt \equiv 1$ and $b$, $\delta \in [0, 1]$ so to interpret rates as probabilities. F) CPIM Markov chain with all transition probabilities.

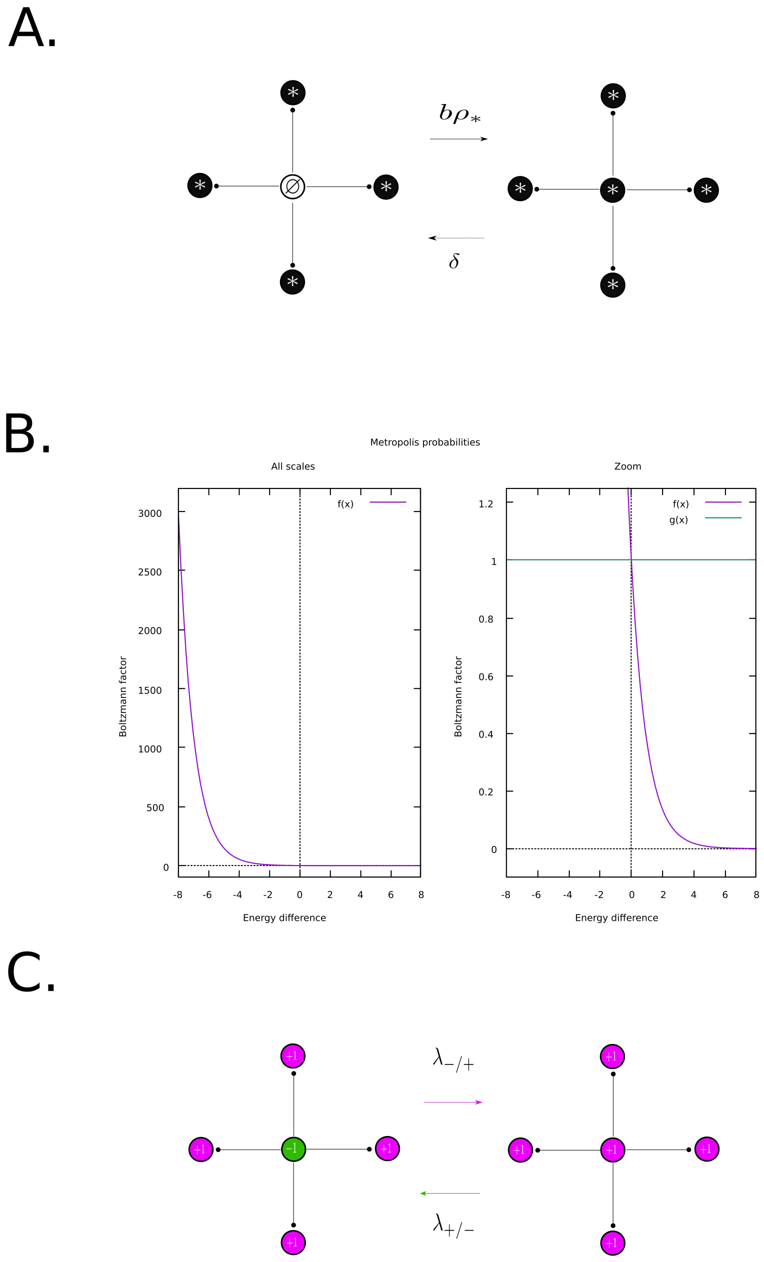

Figure S17: **Neighborhoods and Probability.** A) Nearest Neighborhood configuration with a vacant (focal) site with fully occupied surroundings, which can be colonized at rate $b$ as $\rho_* = 1$, or a reverse situation where a fully surrounded occupied site goes extinct at rate $\delta$. B) Boltzmann factor as a function of energy difference for a proposed flip of a spin. C) Nearest Neighborhood configuration with a single spin down state fully surrounded by spin up states, and with forward spin flip probabilities being $\lambda_{-/+}$ and $\lambda_{+/-}$ for a backwards flip.

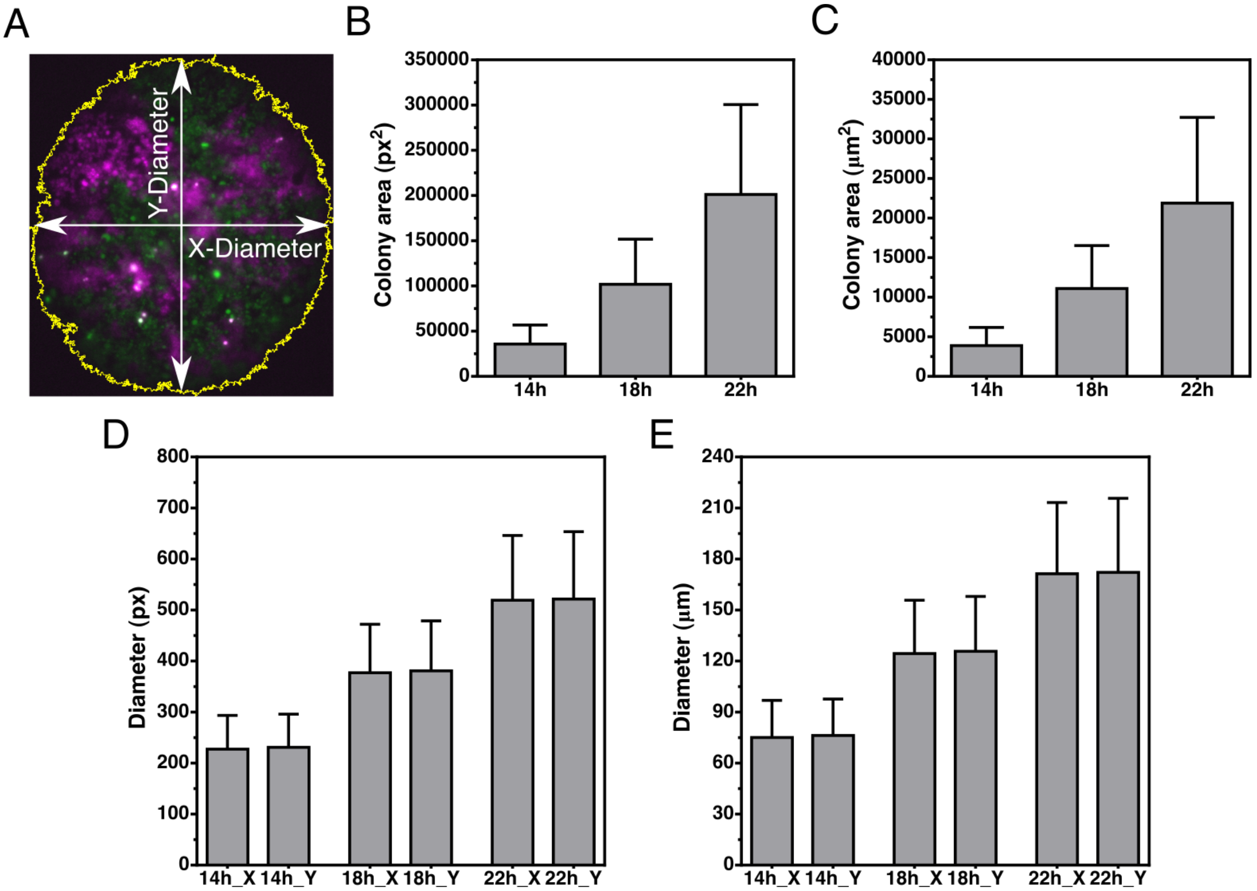

Figure S18: **Colony size expansion in a 3-point time series.** A) Example of a colony and the quantification of its area, x-Diameter, and y-Diameter. B-C) Average size of the colonies in square pixels (B) or square micrometers (C) at 14h, 18h, and 22h post-inoculation. D-E) Average x-Diameter and y-Diameter of the colonies in pixels (D) or micrometers (E) at 14, 18, and 22 hours after inoculation on solid M9-glucose medium. Values and error bars correspond to the mean ± the standard deviation of data obtained from 13 colonies.

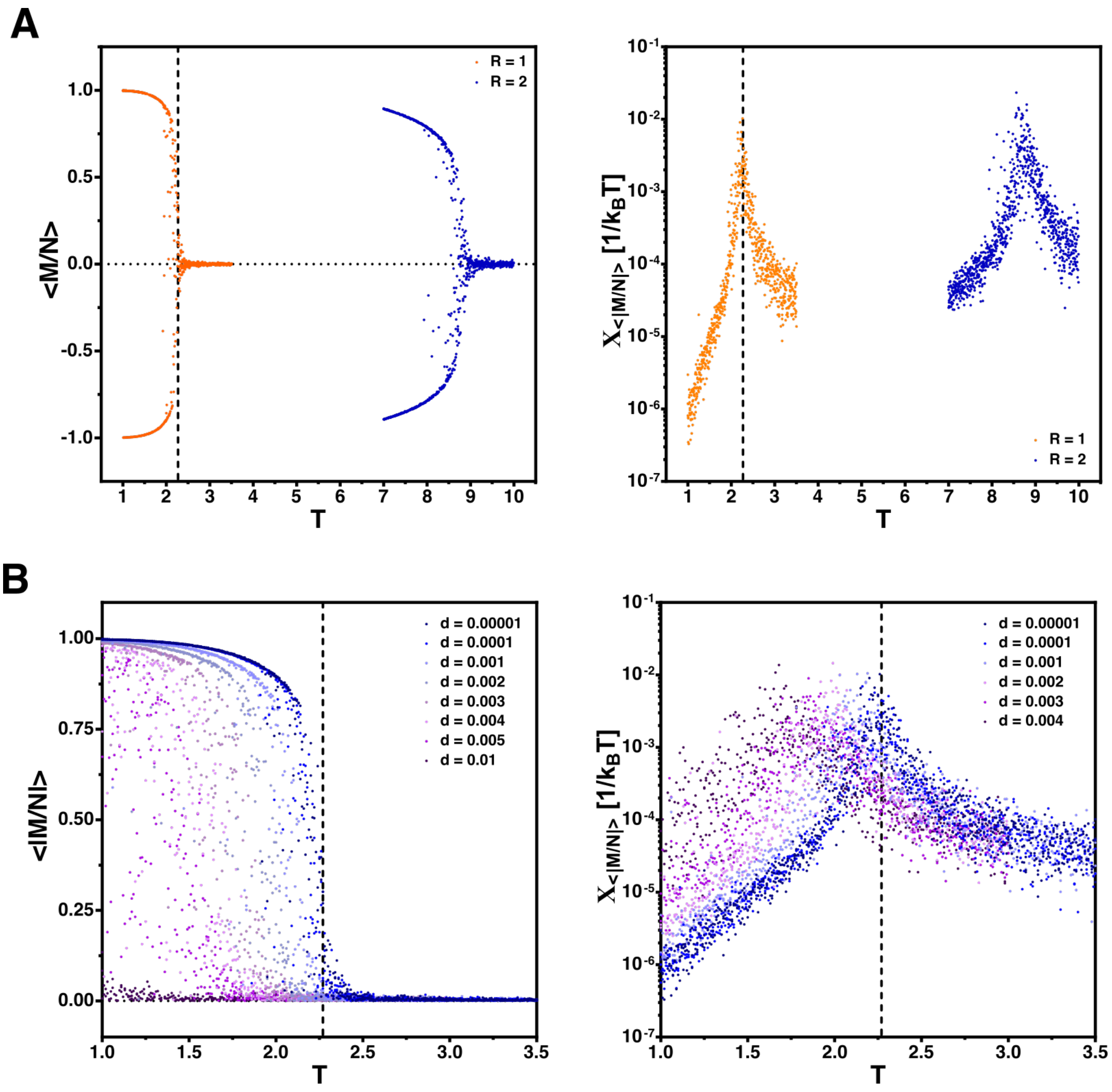

Figure S19: **Behavior of the critical value of $T$ for different parameters of CPIM.** A) Time-averaged magnetization per site $< M/N >$ (left) and magnetic susceptibility per site $X_{<|M/N|>}$ (right) as a function of $T$ for ferromagnetic populations simulated with CPIM for two neighborhoods (R = 1: nearest neighbors, R = 2: next nearest neighbors). For these simulations the following parameters were used: birth rate = 0.03, death rate = 0.00001, differentiation rate = 0.1, lattice = 256x256. 3 simulations per each value of $T$ were used, and to obtain the time average of the magnetization per site, 11 values were taken between generations 6800 and 9800. (B) Absolute values of the time-averaged magnetization per site $< |M/N > |$ (left) and magnetic susceptibility per site $X_{<|M/N|>}$ (right) as a function of $T$ for ferromagnetic populations simulated with the CPIM with nearest neighbors at different values of death rate $d$. Birth rate = 0.03, differentiation rate = 0.1, lattice = 256x256. Dotted vertical lines mark the critical value of $T$ of the Ising model ($T_c = 2.27$).

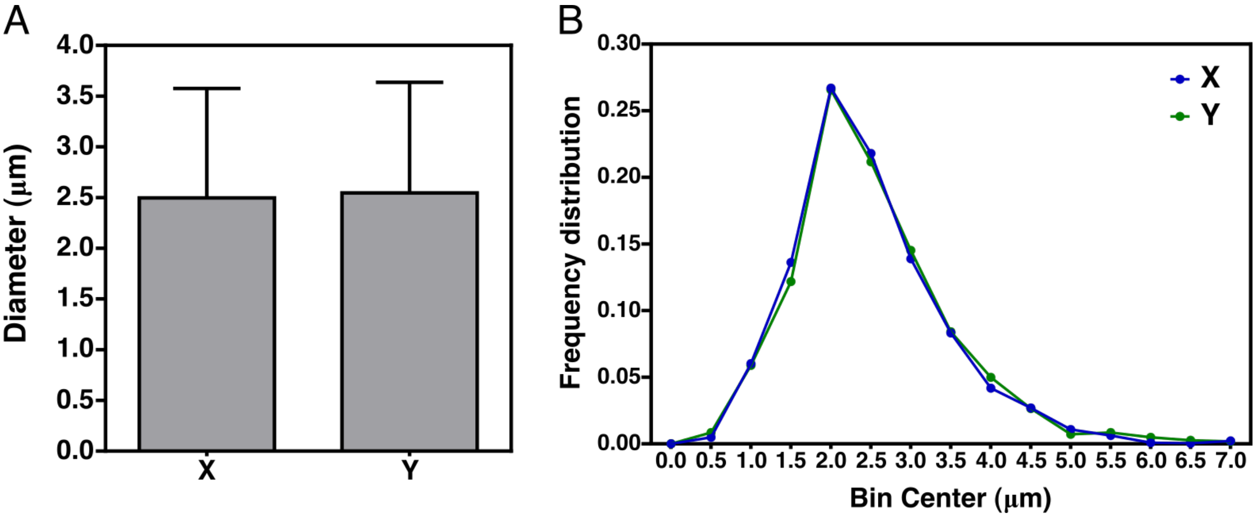

Figure S20: **Average size of spherical *E. coli* cells.** A ferromagnetic colony of the *E. coli* KJB24 strain was re-suspended in M9-glucose medium. This strain contains a mutation in the cell wall protein RodA, generating spherical cells. Cells were observed under the microscope and the average (A) and frequency distribution (B) of the x- and y-diameter of the cells were calculated using the Analyze particles command of the Fiji distribution of ImageJ Schindelin *et al* (2012). Values and error bars in A correspond to the mean $\pm$ the standard deviation of data obtained from 2225 cells.

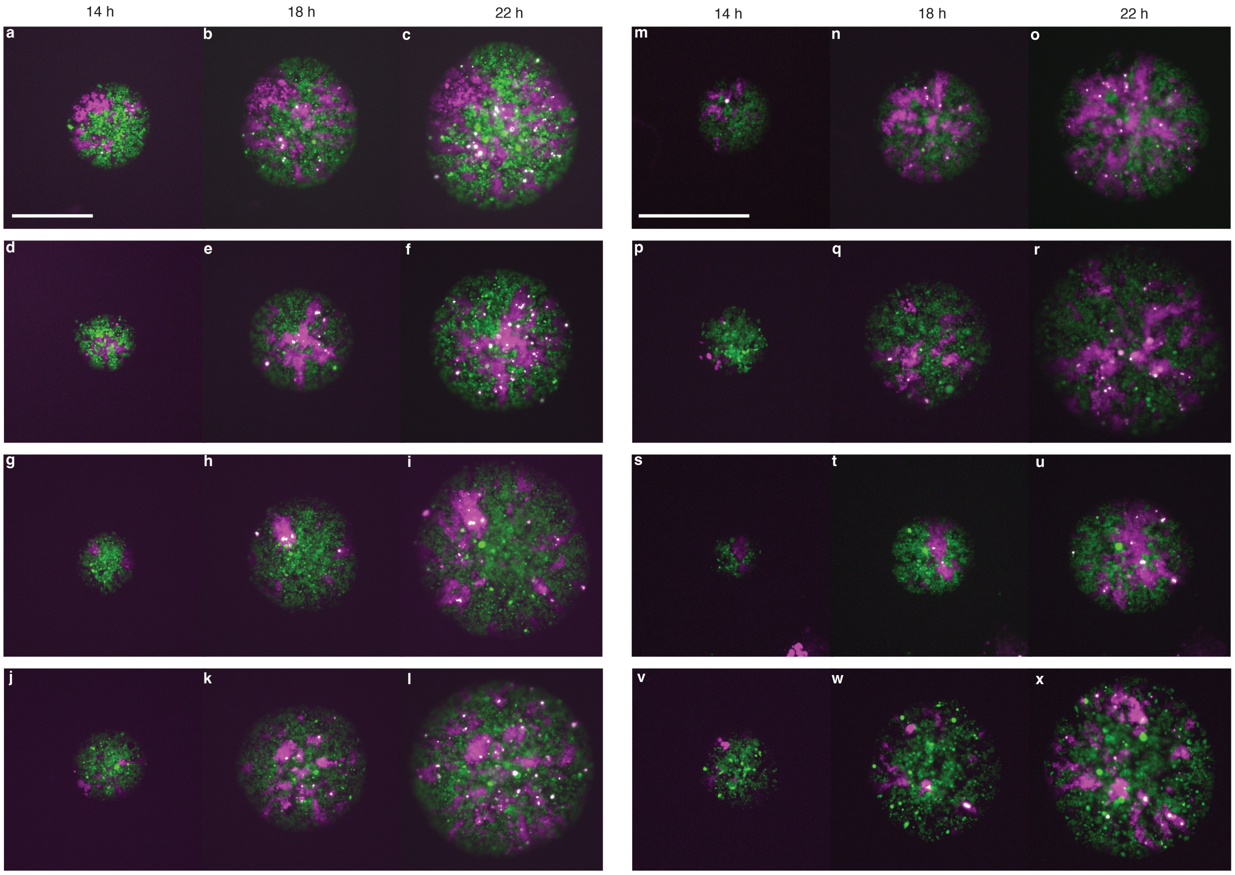

Figure S21: **Time series of colonies of spherical *E. coli* cells carrying the ferromagnetic system with reporter vector 1.** Cells were grown on solid M9-glucose medium supplemented with $10^{-8}$ M of C6HSL, and images were taken 14, 18, and 22 hours after inoculation on the medium. Scale bars 100 $\mu$m.

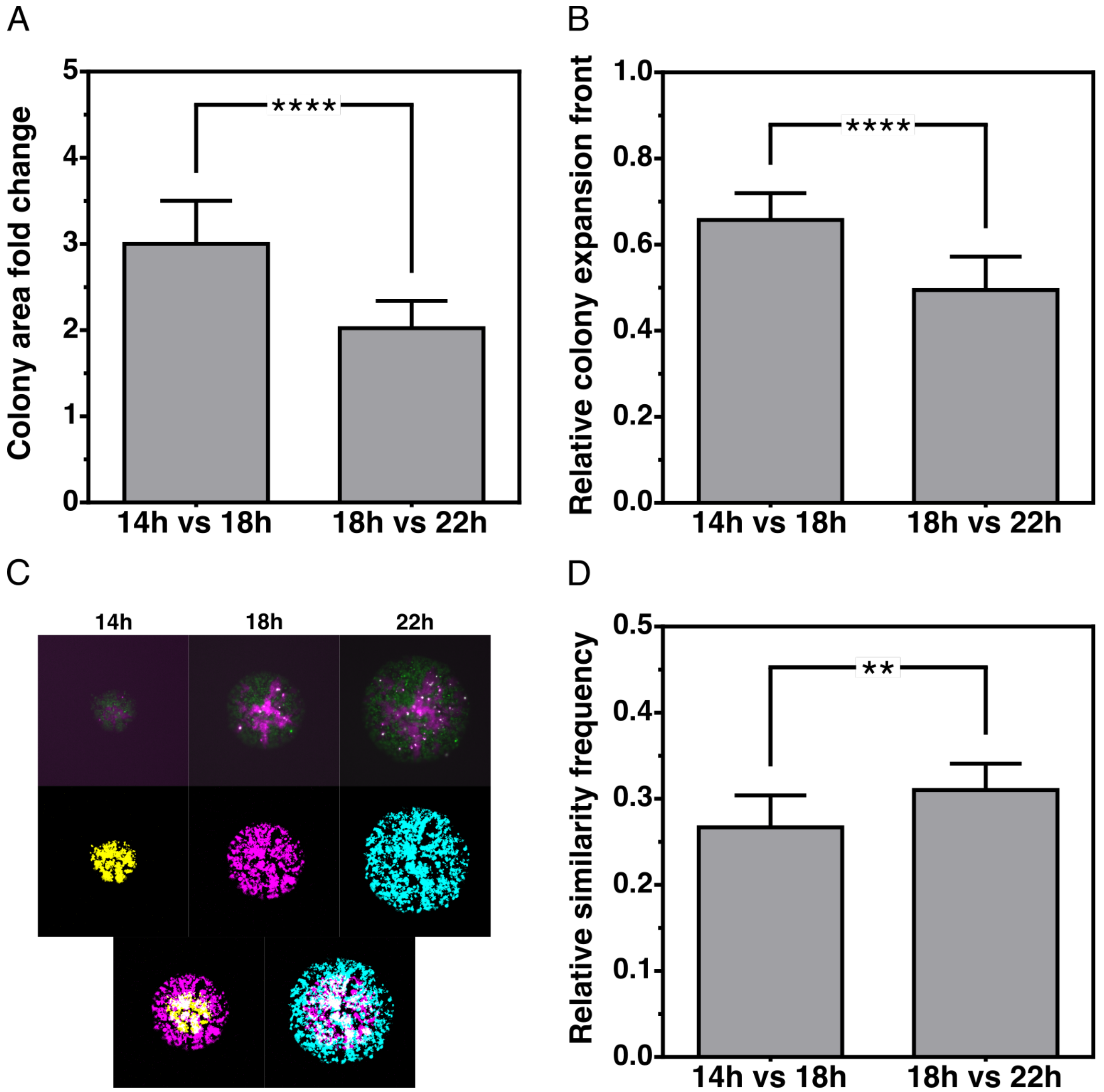

Figure S22: **Dynamics of reporter gene expression (spin flipping rates) in ferromagnetic colonies of spherical *E. coli* cells.** A) Increase of the size of the colonies between 14 and 18 hours, and btween 18 and 22 hours. To calculate the colony area fold change the area of the colonies at 18 (22) hours was divided by the area of the colonies at 14 (18) hours. B) Increase in the size of the expansion front of the colonies between 14 and 18 hours, and 18 and 22 hours. The relative colony expansion front corresponds to the area of the colonies at 18 (22) hours minus the area of the colonies at 14 (18) hours, and divided by the area of the colonies at 18 (22) hours. C) Example of the analysis of the temporal evolution of the patterns that emerge in the colonies. Images were taken at 14, 18, and 22 hours post-inoculation in M9-glucose medium supplemented with $10^{-8}$ M of C6HSL. Top: Merge of the red and green channels, Mid: Binarization of the green channel (for comparison, 14 hours is represented in yellow, 18 hours in magenta, and 22 hours in cyan), Bottom: Merge of the green channels at 18h and 22h, and 14h and 18h. D) Change in reporter expression. Quantification of the similarity between patterns observed at 14 and 18 hours, and 18 and 22 hours. The relative similarity frequency corresponds to the white pixels in C) (where cyan = magenta or magenta = cyan) divided by the total non-black pixels: white/(cyan + magenta + white) or white/(yellow + magenta + white). 13 colonies were used for the analysis, and statistical analysis was performed using unpaired two-tailed Mann-Whitney test ($\alpha = 5\%$). $**$: P value = 0.0019, $****$: $P \leq 0.0001$.

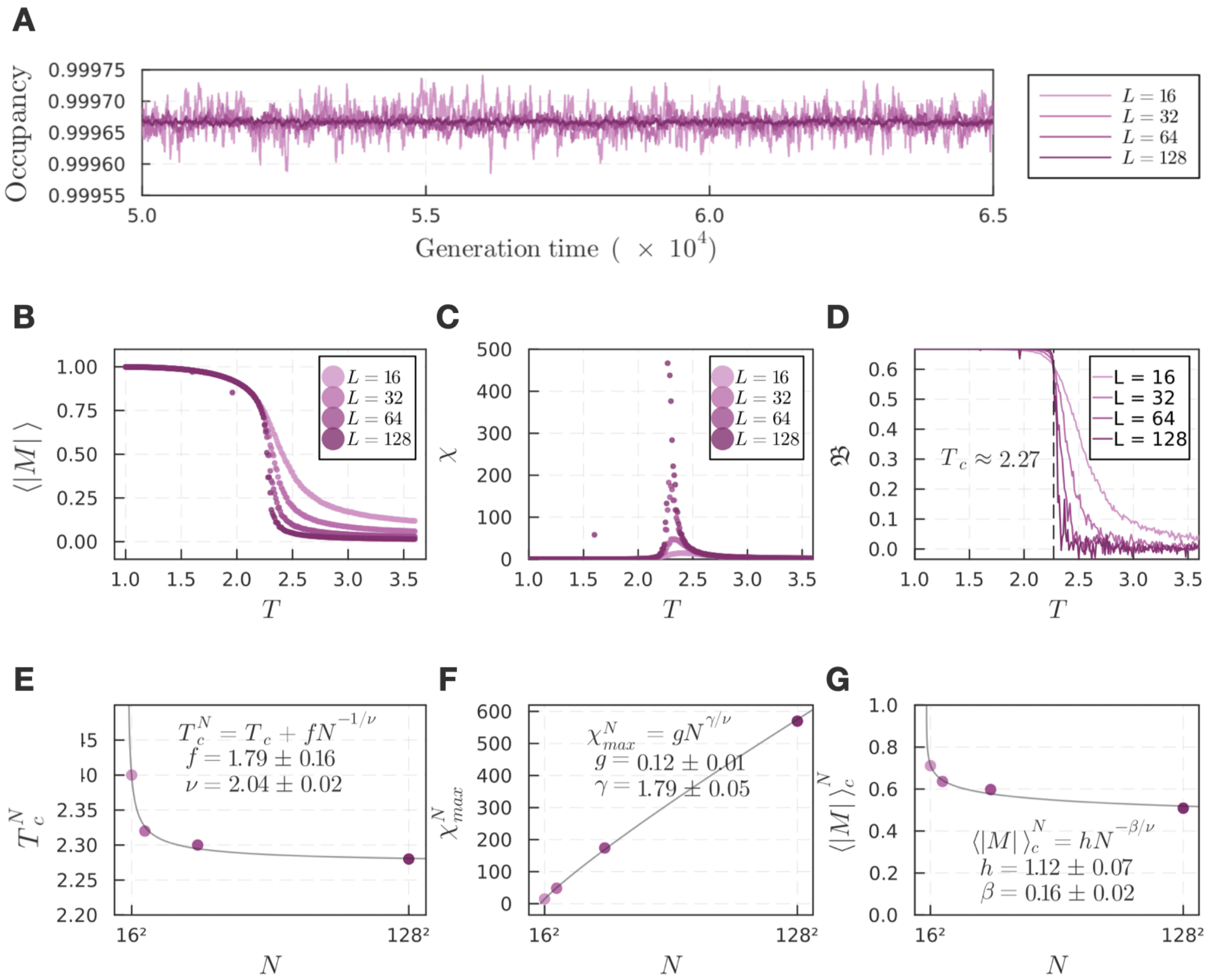

Figure S23: **Finite-size scaling analysis of Contact Process Ising Model (CPIM)** (A) Lattice occupancy $1 - \varnothing/L^2$ (i.e. the density of occupied sites in the lattice) fluctuations over the generation times considered in this study, (B) average magnetization per site $\langle |M| \rangle$, (C) magnetic susceptibility per site $\chi$, and (D) Binder cumulant $\mathfrak{B}$ vs temperature $T$ for different lattice sizes $L = \{16, 32, 64, 128\}$. (E) Position of the magnetic susceptibility per site peak, $T_c^N$, (F) height of the magnetic susceptibility per site peak, $X_{max}^N$, and (G) average magnetization per site at the critical point $\langle |M| \rangle_c^N$ as a function of the system size $N = L^2$. All data points are averages over 15,000 generations and 10 simulations (replicates) using the model parameters $b = 0.03$, $d = 0.00001$, and $\alpha = 0.1$.

Table S1: Plasmids used in this study

| **Name** | **Resistance** | **Ori** | **Relevant features** | **Reference** |
|---|---|---|---|---|
| AFM_1XpSC101 | Kanamycin | pSC101 | Antiferromagnetic vector | This work |
| AFM_1XpSC101_spect | Spectinomycin | pSC101 | Antiferromagnetic vector | This work |
| FM_1XpSC101 | Kanamycin | pSC101 | Ferromagnetic vector | This work |
| FM_1XpSC101_spect | Spectinomycin | pSC101 | Ferromagnetic vector | This work |
| R1_A0_col2 | Chloramphenicol and Carbenicillin | pDestBAC | Reporter vector 1 | This work |
| R2_A0_col2 | Chloramphenicol and Carbenicillin | pDestBAC | Reporter vector 2 | This work |
| SEG10 | Chloramphenicol and Carbenicillin | pDestBAC | SEG vector expressing sfGFP | Nunez *et al* (2017) |
| SEG11 | Chloramphenicol and Tetracycline | pDestBAC | SEG vector expressing mCherry | Nunez *et al* (2017) |

Table S2: Reporter

| | C6HSL | | C12HSL | |
|---|---|---|---|---|
| | *RFP* promoter | *GFP* promoter | *RFP* promoter | *GFP* promoter |
| n | 1.00 | 1.00 | 1.00 | 1.00 |
| $\Delta E$ [$k_B$T] | -3.40 | -4.20 | -3.80 | -4.00 |
| Kdoff [M] | 1.60E-05 | 2.20E-07 | 1.80E-07 | 1.00E-04 |
| Kdon [M] | 4.30E-10 | 1.70E-08 | 6.20E-08 | 2.00E-09 |

Table S3: Ferromagnetic

| | C6HSL | | C12HSL | |
|---|---|---|---|---|
| | *RFP* promoter | *GFP* promoter | *RFP* promoter | *GFP* promoter |
| n | 1.45 | 2.50 | 1.00 | 1.00 |
| $\Delta E$ [$k_B$T] | -6.60 | 6.20 | -6.60 | 5.60 |
| Kdoff [M] | 1.00E-06 | 1.00E-11 | 4.50E-04 | 1.60E-09 |
| Kdon [M] | 1.90E-11 | 2.70E-08 | 1.20E-08 | 2.10E-07 |

Table S4: Anti-Ferromagnetic

| | C6HSL | | C12HSL | |
|---|---|---|---|---|
| | *RFP* promoter | *GFP* promoter | *RFP* promoter | *GFP* promoter |
| n | 1.00 | 1.00 | 1.00 | 1.00 |
| $\Delta E$ [$k_B$T] | -1.20 | -0.40 | -1.20 | -0.30 |
| Kdoff [M] | 9.00E-04 | 1.60E-08 | 1.80E-09 | 1.00E-05 |
| Kdon [M] | 1.80E-08 | 1.40E-07 | 2.60E-08 | 1.80E-09 |

Biological parameters that characterize the binding of the inducers C6HSL and C12HSL to the *RFP* and *GFP* promoters in cells carrying the reporter vector (Table S2), the ferromagnetic system (Table S3) or the anti-ferromagnetic system (Table S4). The values were obtained from the fit to Eq. 2 of the data of red and green fluorescent protein synthesis rates of liquid cultures of cells carrying these systems as a function of the inducer concentrations.

Table S5: Scaling exponent $\gamma$ of populations with ferromagnetic interactions.

| | Scaling exponent $\gamma$ | |
|---|---|---|
| | r_plfit | Least squares fit (inset) |
| CPIM | 1.9413 | 1.989 |
| Rod-shaped Ferro R.V.1 | 2.1739 | 2.175 |
| Rod-shaped Ferro R.V.2 | 2.1369 | 2.086 |
| Spherical Ferro R.V.1 | 1.9095 | 2.126 |
| Spherical Ferro R.V.2 | 1.7593 | 1.866 |
| Spherical Ferro GLY | 2.3073 | 2.227 |
| Spherical Ferro GLU | 2.1918 | 2.141 |

Power-law exponents of ferromagnetic populations simulated with CPIM at $T = T_c$, ferromagnetic colonies of rod-shaped and spherical cells (Fig. 4d), and ferromagnetic colonies of spherical cells grown in minimal medium supplemented with glucose (GLU) or glycine (GLY) (Fig. 4e). The exponents were estimated using the r_plfit(k,'hist') algorithm Hanel *et al* (2017) or by the the least squares method.

Table S6: Scaling exponent $\gamma$ of simulated populations with ferromagnetic interactions at different birth rates.

| | Scaling exponent $\gamma$ | |
|---|---|---|
| Birth rate | r_plfit | Least squares fit |
| 0.0350 | 1.9395 | 1.975 |
| 0.0325 | 1.9381 | 1.983 |
| 0.0300 | 1.9400 | 1.979 |
| 0.0275 | 1.9313 | 1.978 |
| 0.0250 | 1.9244 | 1.962 |
| 0.0225 | 1.9173 | 1.976 |
| 0.0200 | 1.9243 | 2.004 |
| 0.0175 | 1.9262 | 1.974 |
| 0.0150 | 1.9030 | 2.006 |

Power-law exponents of ferromagnetic population of Fig. 4e simulated with CPIM at different cell birth rates. The exponents were estimated using the r_plfit(k,'hist') algorithm (Hanel *et al* (2017)) or by the least squares method.

Table S7: **Numerical critical exponents and critical temperature of the two-dimensional Ising model, Weber-Buceta model Weber and Buceta (2016), and the Contact Process Ising Model (CPIM).**

| Exponent | 2-D Ising model | WB | CPIM |
|---|---|---|---|
| $\nu$ | $0.99 \pm 0.10$ | $1.7 \pm 0.2$ | $2.04 \pm 0.02$ |
| $\gamma$ | $1.78 \pm 0.22$ | $0.8 \pm 0.1$ | $1.79 \pm 0.05$ |
| $\beta$ | $0.13 \pm 0.01$ | $0.1 \pm 0.4$ | $0.16 \pm 0.02$ |
| $T_c$ | $2.27 \pm 0.00$ | - | $2.27 \pm 0.01$ |

Data of the Weber-Buceta model and the 2-D Ising model was taken from Weber M, Buceta J., 2016 Weber and Buceta (2016) and E. Ibarra-García-Padilla *et al.*, 2016 Ibarra-García-Padilla *et al* (2016) respectively.
