## Additional file 3 for "Spatial biology of Ising-like synthetic genetic networks"

July 13, 2023

1. ANID – Millennium Science Initiative Program, Millennium Institute for Integrative Biology (iBio), Santiago, Chile.
2. Institute for Biological and Medical Engineering, Schools of Engineering, Medicine and Biological Sciences, Pontificia Universidad Católica de Chile, Santiago, Chile.
3. Institute of Advanced Studies, Shenzhen X-Institute, Shenzhen, China.
4. Schools of Physics and Biology, Pontificia Universidad Católica de Chile, Santiago, Chile.
5. Department of Natural Sciences and Technology, Universidad de Aysén, Coyhaique, Chile.
6. FONDAP Center for Genome Regulation - Department of Molecular Genetics and Microbiology, Pontificia Universidad Católica de Chile, Santiago, Chile.

Keywords: Ising model, bi-stable, synthetic gene networks, spatial correlation, criticality

### Additional file 3

#### 0.1 Supplementary Note 1: Simulation of the Ising model during the growth of a quasi 2D colony

To study the effects of Ising-like gene-to-gene interactions in a population of bacterial cells growing as a quasi-two-dimensional (2D) colony, we created a lattice model that combines spatial growth dynamics and the Ising model from statistical mechanics to represent the cell state differentiation induced by our genetic network. We combined the simulation of the two-dimensional Ising Model (IM), using the Metropolis algorithm Metropolis *et al* (1953), with the Contact Process (CP) lattice model of spatial stochastic growth Harris (1974) to represent cell population dynamics of the bacterial colonies observed in our range expansion experiments. We name this model CPIM (Contact Process - Ising Model).

CPIM consists of an interacting particle system that follows colonization, extinction process mixed with differentiation dynamics of particles on a two-dimensional lattice ($\mathcal{L}$) of $N$ sites. An important consideration is that colonization/extinction (or birth/death) processes operate at the scale of the entire bacterial cell while differentiation (induction of gene expression) occurs at the scale of molecular networks within living cells. We use this separation of organizational scales to simplify the mixing of the CP and IM.

Each site of this lattice, representing square sectors of the agar surface, can be in one of four states $\mathcal{S} = \{\varnothing, *, -1, +1\}$, which represent vacant locations (state $\varnothing$, white squares), locations occupied with undifferentiated cells (state $*$, black squares), and locations occupied with differentiated cells expressing RFP (state $+1$, magenta squares) or GFP (state $-1$, green squares) reporter genes respectively. The state of a site $x$ at a time $t$ can be described as $\xi_t(x)$ Liggett (2005), an interacting particle system $\xi_t : \mathcal{L} \to \mathcal{S}$ that, for each time point $t$, assign one state from $\mathcal{S}$ to all sites in the lattice. This spatial stochastic process is determined by the following lattice reactions from three different sets of processes: *i*) Birth, death, and differentiation processes: cells die (lattice squares suffer local extinction) and divide at rates $\delta$ and $b$, respectively. For this process an event (local extinction) occurring at rate $\delta$, means that the times $\Delta t_i$ between successive occurrences (lifespan of the occupied states) has an exponential distribution with parameter $\delta$; that is, $\mathcal{P}(\Delta t_i \leq 1) = 1 - \exp(-\delta t)$. Colonization reactions (utilization of vacancy, state $\varnothing$) all depend on the state of nearby sites as they are the sources of colonization propagules. Grouping these states as a super-state $\Omega = \{*, -1, +1\}$ we have that $\xi_t : \mathcal{L} \to \{\varnothing, \Omega\}$ corresponds to a CP. Vacant sites can be colonized only by Nearest Neighbors (NN; $\|x - y\| \leq 1\ \forall x, y \in \mathcal{L}$; *i.e.* north, south, east and west), originating an advancing (2-D) front of growing cells that colonize available habitat in the landscape. This spatial spread, for all $\omega \in \Omega$, is represented by $\varnothing_x + \omega_y \xrightarrow{b\rho_\omega} \omega_x + \omega_y$, while cell death/local extinction is represented by $\omega_x \xrightarrow{\delta} \varnothing_x$. The rate of colonization ($b\rho_\omega$) is proportional to both, the birth rate $b$ and the local density $\rho_\omega$ of $\omega \in \Omega$ states in the NN neighborhood of the focal vacant ($\varnothing_x$) site $x \in \mathcal{L}$.

*ii*) Differentiation and gene expression process. In our experimental system, there is always the possibility that cells do not express our synthetic gene network. We model this as an undifferentiated state $*$ and therefore at the beginning of most simulations, a square representing an undifferentiated cell ($*$ state) is placed in the center of the lattice. Then, we proceed by a Monte Carlo method repeating as many times as there are sites on the lattice to produce one generation (step) of our simulation: a site on the lattice is randomly selected, and if this focal site is vacant ($\varnothing$ state), a random neighbor is selected. If this neighbor is occupied ($\Omega$ states, occurring with probability, $\sum_{\omega \in \Omega} \rho_\omega$), it colonizes the vacant site with probability $b\rho_\omega \cdot dt$. On the other hand, if the focal site is occupied by an undifferentiated state ($*$), if it survives (with probability $1 - \delta \cdot dt$), it can differentiate into a RFP-expressing ($+1$) or a GFP-expressing ($-1$) state with a certain probability $\alpha$, which is represented by $*_x \xrightarrow{\alpha/2} +1_x$ and $*_x \xrightarrow{\alpha/2} -1_x$. As members of the CP's occupancy class $\Omega$, these differentiated states can also die or give birth with rates $\delta$ and $b\rho_*$ respectively. Notice that unlike, birth-death process, internal cell states changes, like genetic differentiation, are conditional upon cell survival, and instead of being determined by rates, they are ruled by probabilities which are conditional on a particle's survival. We use this separation of organizational scales between a complete bacterial cell and a small subset of its constituent components (our synthetic gene network) to make the CP model of vacancy utilization independent from our different distinctions of occupancy ($\Omega$ states).

*iii*) Ising-like cellular states alignment process: as our genetic network consists of a bi-stable switch locally coupled by diffusing small molecules (auto-inducers) that trigger flips between two reporter gene states (GFP and RFP); we model it as spin variables and represent them using the IM (states -1 and +1). To determine if lattice squares representing differentiated cells can change from one state to another (RFP to GFP or GFP to RFP) we use the Metropolis algorithm. Thus, when a site in the lattice which is occupied by a differentiated state (-1 or +1) is randomly selected during a Monte Carlo step, if the square survives extinction (with probability $1 - \delta \cdot dt$) the following computation takes place:

We calculate the interaction energy of the focal site ($i = x \in \mathcal{L}$) and its neighbors ($j = y \in \mathcal{N}$) using $\mathcal{H} = -J \sum_{\langle ij \rangle} \sigma_i \sigma_j$ (Eq. 1; main text). Here, $\sigma_i$ and $\sigma_j \in \{-1, +1\}$ represent spin variables and the notation $\langle ij \rangle$ indicates that the sum is between neighbouring pairs of sites, allowing only a short-range interaction. In most simulations, we considered neighborhoods $\mathcal{N}$ to be NN but we also explored the Next Nearest Neighbors (NNN) case ($\|x - y\| \leq 2 \ \forall x, y \in \mathcal{L}$; where $\| \cdot \|$ is the Manhattan metric). Next, we determine the change in the interaction energy $\Delta\mathcal{H}$, which results if the spin state at that site is flipped. For the interaction energy calculations, we assumed that $J = +k_B$ (where $k_B$ is Boltzmann constant) for ferromagnetic systems and $J = -1k_B$ for anti-ferromagnetic systems (see genetic constructs; main text). This assumption implies that the interaction energy between neighboring sites is minimized when they have the same state in a ferromagnetic system, or the opposite state in an anti-ferromagnetic system (Eq. 1; main text). Thus, if $\Delta\mathcal{H} < 0$, the change in the spin state ($[-/+]_x \xrightarrow{\mathcal{P}} [+/-]_x$) is always accepted (probability $\mathcal{P} = 1$) since this change decreases the total energy of the system. If $\Delta\mathcal{H} \geq 0$, which means an increase in the total energy of the system, the change in the spin state is accepted only with probability $\mathcal{P} = e^{-\Delta\mathcal{H}/k_B T}$ s Landau and Binder (2014). In the classical IM, $T$ is the absolute temperature and corresponds to the control parameter of the system. This means that varying $T$ determines the transition between ordered and disordered configurations of the system. In our model, $T$ represents a parameter that determines the coupling strength between cells in the experimental system: a small value of $T$ represents a strong coupling, while a large value represents a weak coupling.

**Simulations of the model**

In this model, one generation time corresponds to the moment when all the sites in the lattice, statistically all have had the possibility of being visited on average. In a 256x256 lattice, we run simulations for 6800 to 9800 generation times.

Without loss of generality, we consider $k_B = 1$.

**Considerations of our Model assumptions**

It is also important to note that a lattice site in the CPIM simulations can have two interpretations. As described above, a lattice site in the CPIM simulations can represent a cell. However, a lattice site in this model can also represent a cell territory. Considering lattice sites as cell territories is particularly relevant since it allows to map a lattice site to a sector in a colony. In this interpretation, besides considering lattice sites as cell territories (and not individual cells), the death rate represents the clearance rate of territories, and the division/birth rate represents the local colonization rate. However, the usefulness of the model remains the same: local interactions can give rise to large-scale order differences.

**Temporal considerations** The CP has rates defined by the Poisson distribution. In the Mean Field (MF), the proportion of occupied sites, $p$, can be modeled as, $\dot{p} = bp(1-p) - \delta p$ and at equilibrium we have $\hat{p} = 1 - \delta/b$. Thus to have the system persist we need $b > \delta$ and we have a residual vacancy which corresponds to the time units, $b^{-1}$, it takes particles to replicate times their dead/extinction rate. In the case of our colonies, local extinction (death) is negligible at the scale of our observations and compared with the colony expansion rate. Thus in most simulations we used $dt \equiv 1$ with $b = 0.03$ and $\delta = 0.00001$. This means, on average, it takes 100 generations in our simulations for 3 particles to replicate when provided with adjacent vacancy and 100000 generations for them to cease to exist. This mostly takes place at the expansion front of the simulated colonies. The CP representing our colonies is very dense ($\approx 99,9\%$ occupancy. Additional file 2, Fig. S23A) as $b/\delta >> 1$. During this timescale, one generation ($dt$) we decide to use the maximum possible rate for the spin flips of the IM.

If we think on infinite temperature, $T \to \infty$, we have that in one generation (Monte Carlo step; $dt \equiv 1$), all spins flip on average. On the contrary, at zero temperature, spins flip with probability one if the energy difference is negative and not at all if it is positive. When the temperature is small spins always flip for negative energy difference and with exponentially decaying probabilities when energy differences are positive.

The important quantity when simulating the IM is the energy difference

$$\Delta\mathcal{H} = -2J[N_+ - N_-] \cdot \xi_t(x)$$

obtained by flipping a state $\xi_t(x) \in \{-1, +1\}$ to its opposite sign. Here $N_+$ and $N_-$ corresponds to the number of sites with spin states $+1$ and $-1$ in the neighborhood $\mathcal{N}$ of $x$ respectively. The Metropolis algorithm accepts a spin flip with probability $\lambda_{\text{flip}}$ depending on the energy difference between states before and after the proposed flip (see Additional file 2, Fig. S17, B and C). This probability is not normalized by the partition function and all configurations with $\Delta\mathcal{H} \leq 0$ are flipped within a Monte Carlo step ($\lambda_{\text{flip}} = 1$). On the contrary, if $\Delta\mathcal{H} > 0$ then we flip with probability

$$\lambda_{\textbf{flip}} = e^{-\Delta\mathcal{H}/k_B T} = e^{2\cdot\xi_t(x)[N_+ - N_-]/T}. \quad (1)$$

Notice that the Markov process produced by a Monte Carlo step on the lattice (see Additional file 2, Fig. S16C) is not meant to represent flipping rates but rather to draw configurations that are consistent with the Boltzmann distribution given by the Hamiltonian of Eq.1 (main text). Therefore in the CPIM, energy-favorable flips are considered to be an instantaneous process that is conditional to CP's particle survival. Without loss of generality, as it only introduces transient states we include an undifferentiated state which turns into spins (differentiated states) with no bias with probability $\alpha = 0.1$. Thus in one generation of the CPIM, 10% of surviving undifferentiated occupied sites, become spin variables of the IM.

**Majority rules** The CPIM has transition probabilities that depend on the configurations of neighborhoods. The colonization of an empty site in the CP occurs at rate $b$ when the vacant site is fully surrounded by occupied ones (or probability $b \cdot dt$). In Fig. S17A (Additional file 2) we see such a vacant site fully surrounded ($\rho_* = 1$) by the undifferentiated state $*$. On the contrary, sites can go extinct independently of neighborhood configuration at rate $\delta$ (or probability $\delta \cdot dt$). When making distinctions of occupancy states ($\Omega$), their neighborhood densities $\rho_{\omega\in\Omega}$ determines their colonization likelihood. For the IM, there are a finite number of neighborhood configurations of sites that are in spin states. As we see in Fig. S17, B and C (Additional file 2) and Equation 1 there is an even smaller number of possible spin occupancy patterns $(N_-, N_+)$ for these neighborhoods. For the case of NN interactions, if Temperature is $T = 1$ and $J = k_B^{-1}$, we only have 11 possible values of $\Delta\mathcal{H}$ and their corresponding flipping probabilities.

| NN Neighborhood configurations at $T = 1$ & $J = k_B^{-1}$ | | | | | |
|---|---|---|---|---|---|
| $\Delta\mathcal{H}$ | $e^{-\Delta\mathcal{H}}$ | $\lambda_{\text{flip}}$ | $\Delta\mathcal{H}$ | $e^{-\Delta\mathcal{H}}$ | $\lambda_{\text{flip}}$ |
| -8 | 2980.95 | 1 | +1 | 0.36787 | 0.36787 |
| -6 | 403.428 | 1 | +2 | 0.13533 | 0.13533 |
| -4 | 54.5981 | 1 | +4 | 0.01831 | 0.01831 |
| -2 | 7.38905 | 1 | +6 | 0.00247 | 0.00247 |
| -1 | 2.71828 | 1 | +8 | 0.00033 | 0.00033 |
| 0 | 1 | 1 | | | |

That is, even at small temperature values, all spin-flips with favorable energy are executed within the time scale of one generation on all surviving particles. As we do not have detailed information on the diffusion of HSL molecules in the agar, nor on their transport rates we decide to use the IM as implemented by the Metropolis algorithm when hybridizing it with the CP. Most likely configurations with higher energy differences in favor of a flip would happen at a higher rate, the large order of magnitudes in differences will introduce numerical complexity when running simulations. Thus, as a first approach, we decide to leave the IM as it's implemented in the Metropolis algorithm and use the flipping pseudo rates as rates of our model. In practice, this means that a few neighboring sites favoring a given flip will be enough to trigger it. If vacancy is abundant, however, the CP alters the IM as persistent isolated sites (surrounded by vacancy) have zero interaction energy independent of their spin value and are will flip with probability one. Frustrated sites where the spin occupancy $N_+ - N_- = 0$ also flip instantaneously. This reflects the bi-stable nature of genetic network and our ignorance of its characteristic time-scales (see Section 0.3 hereunder)

#### 0.2 Supplementary Note 2: Finite-size scaling analysis

Studying the effects of the finite size of the system on observables that characterize the model's Ising-like critical behavior such as the average absolute value of the magnetization per site $\langle |M| \rangle$ (Additional file 2, Fig. S23B), and consequently, the magnetic susceptibility per site $\chi = N(\langle |M|^2 \rangle - \langle |M| \rangle^2)$ (Additional file 2, Fig. S23C), allows us to assess whether the phase transition exhibited in the CPIM falls under the universality class of that of the Ising model. This can be done by determining the critical exponents $\nu$, $\gamma$, and $\beta$ (Additional file 2, Fig. S23, E, F, and G respectively; Table A7) as done in Weber M, Buceta J., 2016) Weber and Buceta (2016). This task was carried out by running CPIM simulations for a range of temperatures $T \in [1.0, 3.6]$, on different lattice sizes $L = \{16, 32, 64, 128\}$, using the same model parameters $b = 0.03$, $d = 0.00001$, and $\alpha = 0.1$. All simulations were initialized from a single undifferentiated lattice site, run for $50,000$ generations to ensure the stabilization of the colonization/extinction dynamics (Additional file 2, Fig. S23A), and finally, data was recorded for $15,000$ generations. Each simulation was repeated 10 times (replicates), critical exponents are means over the replicates and their respective standard deviations are shown.

As seen in Fig. S23, B and C (Additional file 2), the critical temperature value $Tc$ at which the (magnetic) phase transition occurs is difficult to pinpoint when the system's size is too small. This problem can be circumvented by means of the cumulant intersection method developed by K. Binder Binder and Landau (1984), which consists of comparing the Binder cumulant $\mathfrak{B}$ (Equation 2) as a function of the temperature $T$ for different lattice sizes $L$ (Additional file 2, Fig. S23D).

$$\mathfrak{B} = 1 - \frac{\langle M^4 \rangle_L}{3\langle M^2 \rangle^2} \tag{2}$$

The intersection of the Binder cumulant $\mathfrak{B}$ curves marks the critical temperature of the system in the thermodynamic limit $T_c \approx 2.27 \pm 0.01$ (Additional file 2, Fig. S23D, dashed line) since $\mathfrak{B}$ does not depend on the system size at the critical point Binder (1981).

After finding the critical temperature $T_c$ at which the system undergoes an Ising-like phase transition, a fit to the finite-size scaling relation

$$T_c^N = T_c + fN^{-1/\nu} \tag{3}$$

was used to determine the value of the critical exponent $\nu$: $\nu = 2.04 \pm 0.02$, where $T_c^N$ corresponds to the position of the peak of the magnetic susceptibility per site for each lattice size studied (Additional file 2, Fig. S23E) and $f$ is a constant.

Furthermore, both the height of the peak of the magnetic susceptibility per site $\chi_{max}^N$ and the average magnetization per site at the critical point $\langle |M| \rangle_c^N$ depend on the system's size following power-laws (Additional file 2, Fig. S23, F, and G respectively), so a fit to

$$\chi_{max}^N = gN^{\gamma/\nu} \tag{4}$$

where $g$ is a constant, yields $\gamma = 1.79 \pm 0.05$. Finally, fitting the simulations data to

$$\langle |M| \rangle_c^N = hN^{-\beta/\nu} \tag{5}$$

where $h$ is a constant, reveals the critical exponent $\beta$: $\beta = 0.16 \pm 0.02$.

All critical exponents calculated in this work are means (of generation time averages) over 10 replicates, and standard deviations of these means are presented as uncertainty measures. The critical exponents calculated here and their respective values for the Weber-Buceta model Weber and Buceta (2016) and a numerical solution of the two-dimensional Ising model are summarized in Table A7).

#### 0.3 Supplementary Note 3: Time series analysis

To gain insight into the dynamics of reporter gene expression patterns in our colonies and how to relate these with the spin flipping rates produced by the Metropolis algorithm of the Ising Model, we compared pictures of experimental colonies taken at tree snapshots separated by 4 hours of growth. Examples of these colonies are shown in Additional file 2, Fig. S21. These correspond to: (i) 14 hours, (ii) 18 hours, and (iii) 22 hours post inoculation respectively. In Fig. S18 (Additional file 2) we show how we can estimate colony expansion by measuring changes in its area and diameter. By comparing colonies between two successive snapshots, we can define a core and an expansion ring as shown in Fig. S22C (Additional file 2). Notice that colony expansion is significantly slower when comparing 18 hours and 22 hours in contrast to what is observed when comparing 14 hours with 18 hours (see Additional file 2, Fig. S22, A, and B). We can estimate the sectors of the colony which change the pattern of expression and observe, as shown in Fig. S22D (Additional file 2), that the core, defined by two consecutive snapshots reflects more similarity in the pattern of gene expression (white pixels) as the colonies grow older. This can be due to entry into stationary phase or other stresses experienced by the colony as its radius becomes larger.

Although our CPIM approach considers a constant and spatially homogeneous rate of spin flipping an interesting avenue for future theory development is to develop models where this can vary in space and time. To be able to develop such models would be extremely important to have experiments that measure the dynamics of these patterns in gene expression under different ecological conditions of food supply and waste removal.

#### 0.4 Supplementary Note 4: Comparison of our work with the work of Weber and Buceta

The work of Weber and Buceta (WB; Weber and Buceta (2016)) is a relevant study to keep in mind when analyzing our experimental results. In their work, they propose a proof of concept model of a gene regulatory circuit. Although our theoretical analysis and simulations differ considerably from their work, both can be used in a complementary fashion to interpret our experiments as both rationals are inspired by the Ising model. The first consideration to have in mind is that our work is guided by experiments and we develop theory to interpret those specific experiments. Secondly, our experiments are on range expansion of an *E. coli* colony growing on a two-dimensional hard agar surface and therefore we focus on a spatially structured model of growth and not on modeling the specifics of the underlying molecular network. However, we do use a complementary model to calibrate the molecular constructs we use for coupling our genetic switch (see Eq. 2 and Fig. 2.; main text). Although in both cases the use of autoinducer molecules is suggested to couple bi-stable switches, our coupling constructs are different from the proof of concept propose in WB. Our experimental genetic switch is also different from the work of Gardner *et al* (2000) which corresponds to the toggle switch WB coupled conceptually in their simulations. Another important difference is that in our work we explored both, ferro and anti-ferro magnetic interactions experimentally. Although our simulations as well as genetic details of our molecular constructs and those of WB are different, we can interpret our experiments while comparing both theoretical approaches (see Discussion; main text).

The switch studied in Gardner *et al* (2000) corresponds to proteins LacI ($U$ in WB) and $\lambda$ ($V$ in WB) which define a concentration pair $(u, v)$ that acts as a toggle switch which in ideal theoretical conditions can be thought to exists in two alternative configurations (phenotypes) $s_1(u_{s_1}, v_{s_1}) = (5.38, 0.508)$nM (spin up, state +1) or $s_2 = (u_{s_2}, v_{s_2}) = (v_{s_1}, u_{s_1})nM$ (spin down, state -1). As they point out, in a volume of $1.5\mu$m$^3$ (typical *E. coli* cell) this corresponds to 0.5 and 5 molecules per cell on average. A number that will be challenging to measure experimentally. Even considering noisy fluctuations with an amplitude reaching 15 molecules per cell (Fig. 2 in WB) would still be a challenging experiment. In our case, we used fluorescent reporters which are expressed in concentrations easily observable by fluorescent microscopy while focusing our attention on a macroscopic colony expanding over a surface of hard agar. WB, on the other hand, examine a different demographic context when they turn their attention to an unstructured theoretical population (Fig. 5 therein, Weber and Buceta (2016)). Spatial structure is of crucial importance in our experimental setup and therefore we use the 2D Contact Process (CP) to model our expanding colony (see main text, and Sections 0.1 and 0.3 of this document).

In their simulations, WB uses an original toggle switch (module 1 in their Figure 2), in which repressors (LacI and $\lambda$cI) are regulated by constitutive-repressible promoters, in parallel to a HSL-regulated expression of receptors (LuxR/LasR) and a second copy of repressors (LacI and $\lambda$cI) (module 2 and 3 in their Figure 2). Therefore, they have HSL-regulated receptors (LuxR/LasR) and two copies of repressors. In our system, we have only one copy of each repressor. Also, the LuxR/LasR are expressed constitutively. The coupling signals in our system regulate the expression of inducer-producing enzymes LuxI and LasI and repressors

LacI and TetR; whereas in WB, HSL regulates one of the two copies of repressors and LuxR/LasR. Thus, we have inducible-repressible promoters that are induced by HSL and repressed by repressors. These promoters regulate the repressors, LuxI/LasI, and the reporters. In our system, repressors (LacI/TetR) and inducers (LuxI/LasI) genes are regulated at the same time by these inducible-repressible promoters (pLux-pTet regulating LacI and LuxI, pLas-pLac regulating TetR and LasI). We use LacI and TetR as repressors, while they use LacI and $\lambda$cI.

WB assumes no cross-talk while we do observe it to exist experimentally at high auto-inducer concentrations. This fact together with the induction of stationary phase limit the time of growth and the size of colonies to be considered in our experiment. Partition of transcription space in *E.coli* means that the polymerase changes its affinity for some promoters at different phases.

WB develop an "effective" approach by focusing on the fact that $U$ and $V$ act as their own auto-inducers. They use this fact to formulate a model which depends on an effective transport parameter $D$ regulating concentrations of actors in intracellular ($U_i$ and $V_i$) as well as extracellular compartments ($U_e$ and $V_e$). Transport between compartments is modeled as $rD$ where $r$ represents the ratio between compartment volumes, $r = V_{cell}/V_{ext}$. A crucial point is their assumption that diffusion across compartments is at least one order of magnitude different than diffusion of auto-inducers in the extracellular compartment. The CPIM does not attempt to model transport or concentrations of proteins as it rather focuses on spatial occupancy. It models the spatial spread of occupancy over vacancy. To model the genetic switch we simply make three distinctions of occupancy types (lattice sites): un-differentiated site (state $*$), spin down (state -1), and spin up (state $+1$). In some sense, our model is also an "effective" theory in the sense that our experimental spatial data (pixels of a fluorescent image) corresponds to area sectors of agar, where autoinducer molecules that have accumulated in both intracellular as well as extracellular space can induce expression of our reporters in that territory. In the case of ferromagnetic experiments a pixels induce the same state on their neighbors and in the case of anti-ferromagnetic case, they induce the opposite state. We represent this by the Ising Hamiltonian and its corresponding sign for parameter J values. Only our ferromagnetic experiments can be interpreted in the light of WB. However, we do not have detailed information about how effective are auto inducers when triggering changes in gene expression, nor their actual concentrations in the agar surface or their diffusion rates. Our model is as generic as possible to be able to be calibrated when further experiments in microfluidic devices are performed.

The other point is that when the reproductive number of the CP is high enough, our model is more similar to the IM than that of WB and our exponents might be even more similar to the pure IM. Just like the IM, in the CPIM, temperature $T$ and coupling $J$ are independent.

WB assume a concentrated culture condition to develop their order parameter. We use the Temperature parameter of the IM as our control parameter.

The most important part is the analogy between Temperature and auto-inducer concentration and transport. This produces a critical value of inducer which triggers the transition.
