## Additional file 1 for "Spatial biology of Ising-like synthetic genetic networks"

July 12, 2023

1. ANID – Millennium Science Initiative Program, Millennium Institute for Integrative Biology (iBio), Santiago, Chile.
2. Institute for Biological and Medical Engineering, Schools of Engineering, Medicine and Biological Sciences, Pontificia Universidad Católica de Chile, Santiago, Chile.
3. Institute of Advanced Studies, Shenzhen X-Institute, Shenzhen, China.
4. Schools of Physics and Biology, Pontificia Universidad Católica de Chile, Santiago, Chile.
5. Department of Natural Sciences and Technology, Universidad de Aysén, Coyhaique, Chile.
6. FONDAP Center for Genome Regulation - Department of Molecular Genetics and Microbiology, Pontificia Universidad Católica de Chile, Santiago, Chile.

Keywords: Ising model, bi-stable, synthetic gene networks, spatial correlation, criticality

### Additional file 1

### Supplementary figures S1-S11

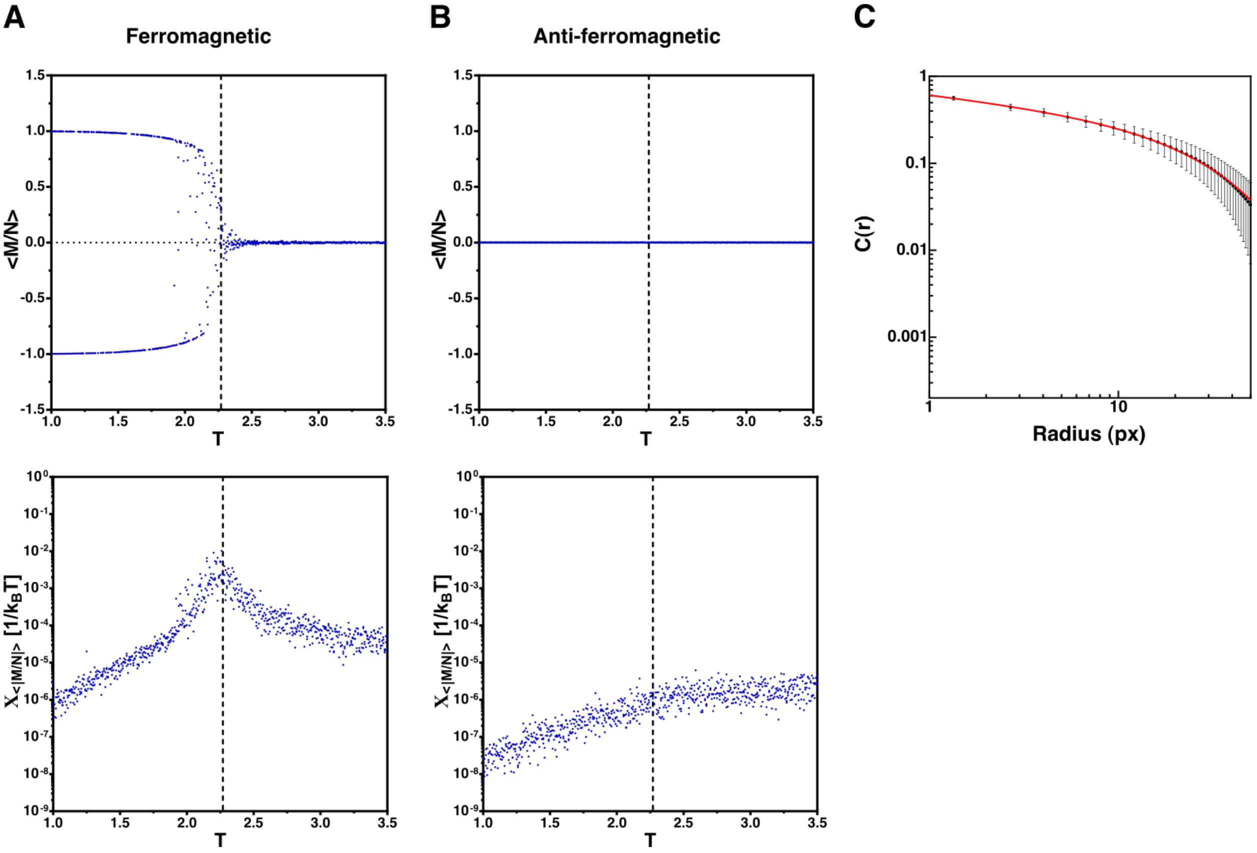

Figure S1: **Power-law decay of the sACF in ferromagnetic populations simulated using CPIM.** Time-averaged magnetization per site $< M/N >$ (top) and magnetic susceptibility per site $X_{<|M/N|>}$ (bottom) as a function of $T$ for ferromagnetic (A) and anti-ferromagnetic (B) populations simulated with the CPIM model. Dotted vertical lines mark the critical value of $T$ ($T_c$) for the Ising model. For these simulations the following parameters were used: birth rate = 0.03, death rate = 0.00001, differentiation rate = 0.1, lattice = 256x256, R = 1 (nearest neighbors). In ferromagnetic populations, below $T_c$ most lattice sites have the same state due to the strong coupling, and the absolute value of $< M/N >$ ($< |M/N| >$) is close to 1. Above $T_c$, lattice sites can randomly adopt a state independent of the state of their neighbors, and $< |M/N| >$ is close to 0. The value of $< M/N >$ for a population with anti-ferromagnetic interactions showed to be always close to zero, independent of the value of $T$. 3 simulations per each value of $T$ were used, and to obtain the time average of the magnetization per site, 11 values were taken between generations 6800 and 9800. (B) sACF of simulated populations with ferromagnetic interactions at the critical value of $T$ ($T = T_c$). The sACF showed to follow a power-law decay with a critical exponent $\eta = 0.2470$. Black dots and error bars correspond to the mean ± the standard deviation of the sACF of 50 simulations at $T = T_c = 2.27$, and the red line correspond to the best fit of the data to the curve $C(r) = A\frac{exp(-r/B))}{r^{\eta}}$, where $\eta$ corresponds to the critical exponent of the autocorrelation function. The radius is in pixels.

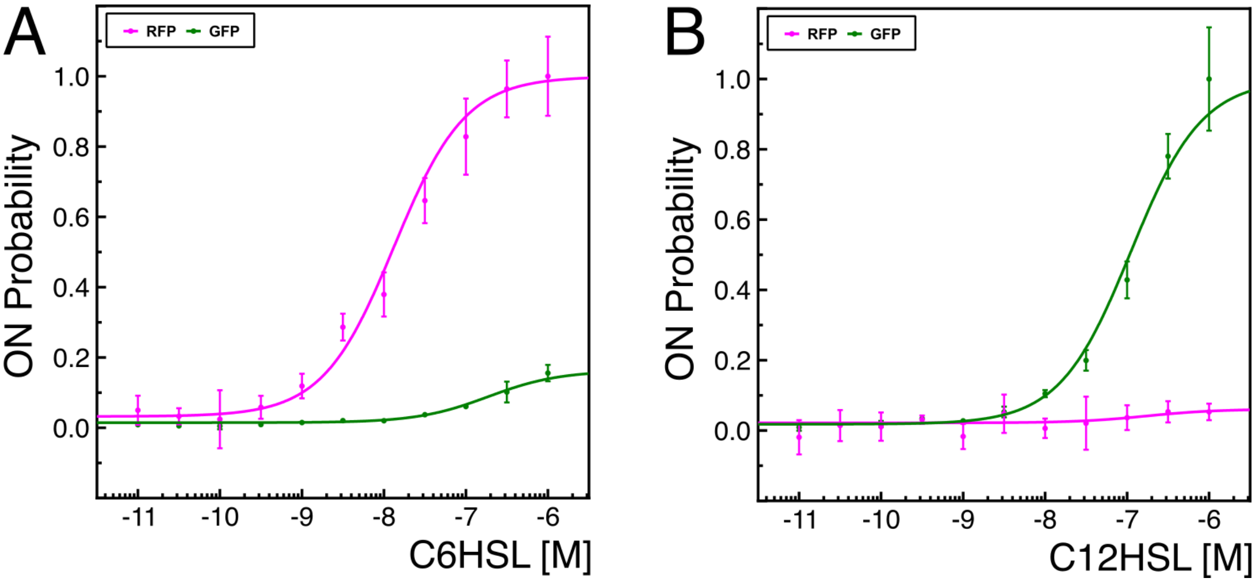

Figure S2: **Red and green fluorescent protein synthesis rate of *E. coli* cells carrying the reporter vector.** Cells were grown in liquid media supplemented with different concentrations of C6HSL (left) and C12HSL (right). Points and error bars correspond to the values of the fluorescent protein synthesis rates normalized by the maximum value reached in each system, and lines correspond to the fit of the data to Eq. 2. Values on the x-axis correspond to the $\log_{10}$ of the C6HSL concentration.

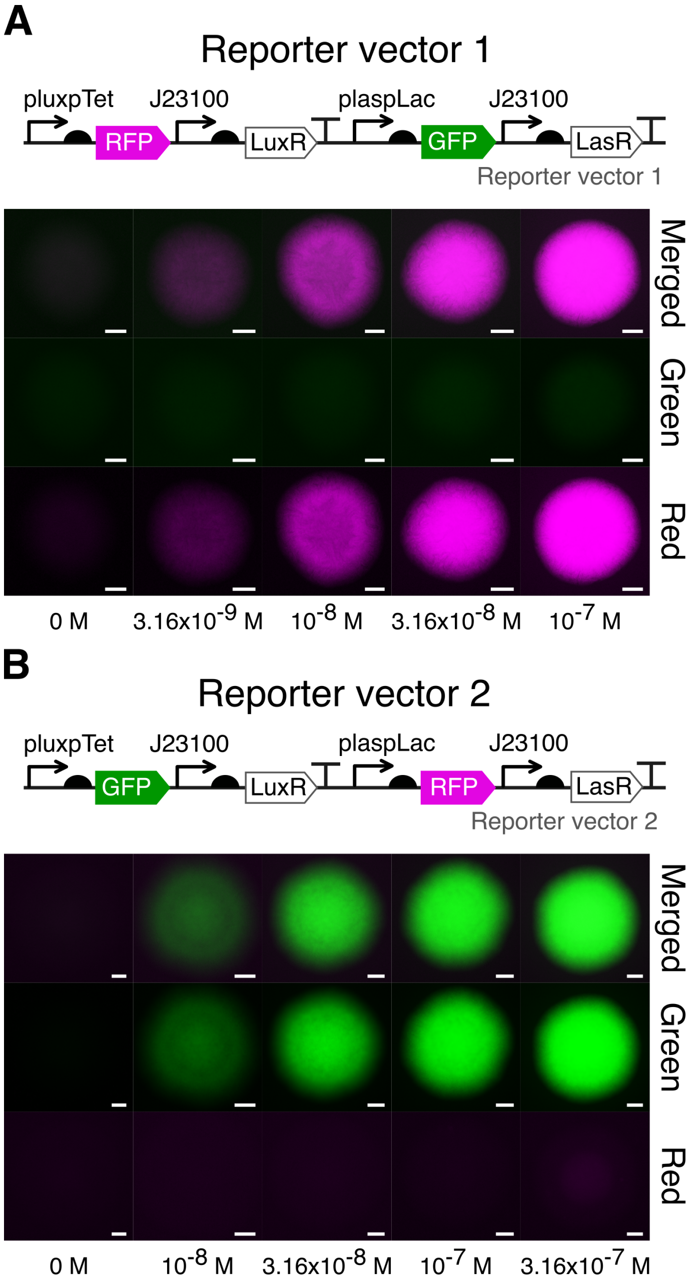

Figure S3: **Induction of red and green fluorescence by C6HSL in colonies of rod-shaped cells carrying different versions of the reporter vector.** Red and green fluorescence of colonies of cells carrying the reporter vector 1 (A) and reporter vector 2 (B) grown on solid M9-glucose media supplemented with different concentrations of C6HSL. Gene network arrangement of reporter vectors 1 and 2 is included for comparison. Images were taken approximately 14 hours after inoculation. Scale bars 100 $\mu$m.

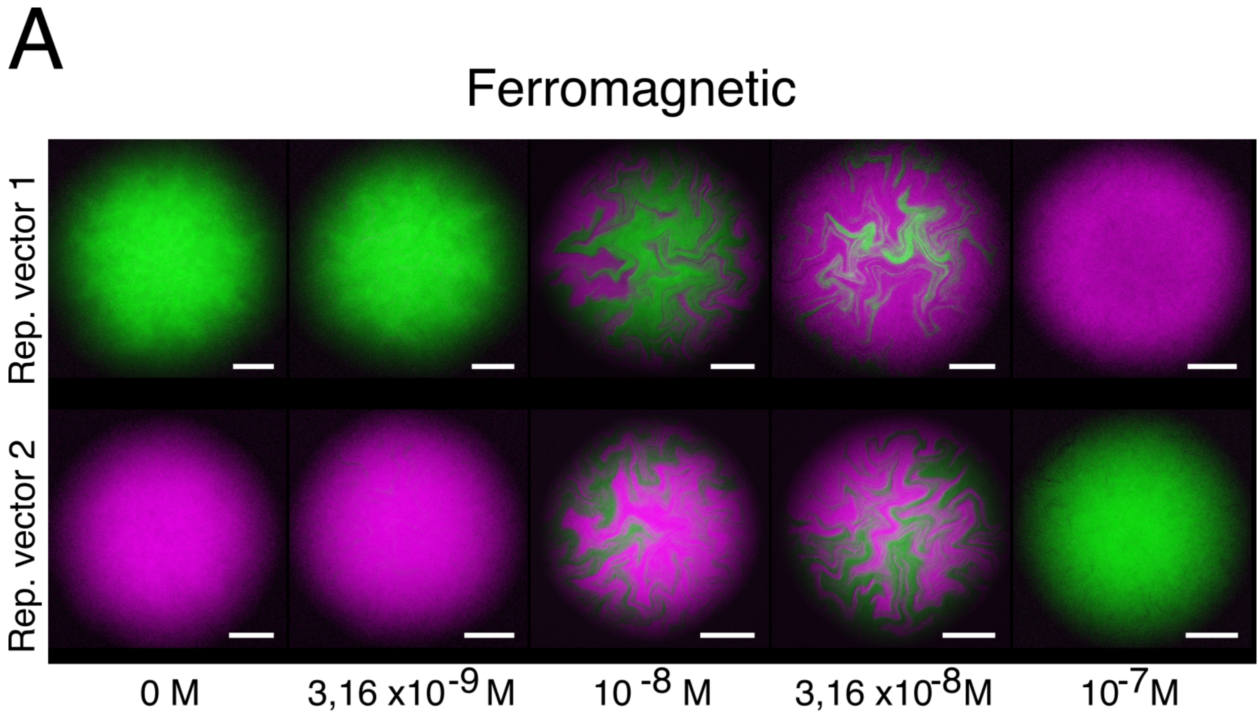

Figure S4: **Red and green state patterns generated in ferromagnetic and anti-ferromagnetic colonies at different concentrations of C6HSL.** Representative images of the cellular state patterns generated in colonies of rod-shaped *E. coli* cells carrying the ferromagnetic (A) or anti-ferromagnetic (B) vector along with the reporter vector 1 or 2. Cells were grown on solid glucose-M9 media supplemented with the indicated concentration of C6HSL. Images were taken approximately 14 hours after inoculation. Scale bars 100 $\mu$m.

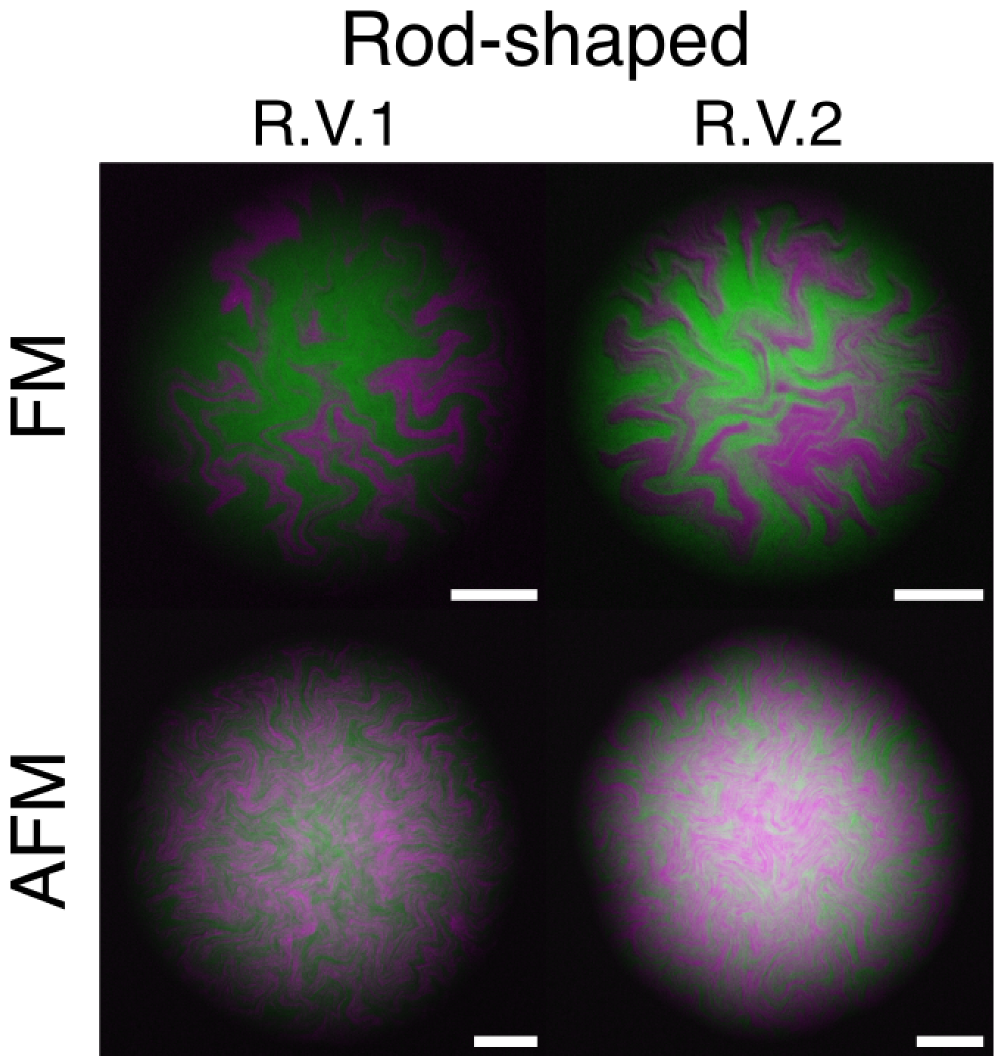

Figure S5: **Cellular state patterns in ferromagnetic and anti-ferromagnetic colonies.** Representative images of red and green fluorescent protein patterns in colonies of rod-shaped *E. coli* cells carrying the ferromagnetic (FM) or anti-ferromagnetic (AFM) systems with reporter vector 1 (R.V.1) or 2 (R.V.2). Cells were grown on solid M9-glucose medium supplemented with $10^{-8}$ M of C6HSL. Images were taken approximately 14 hours after inoculation. Scale bars 100 $\mu$m.

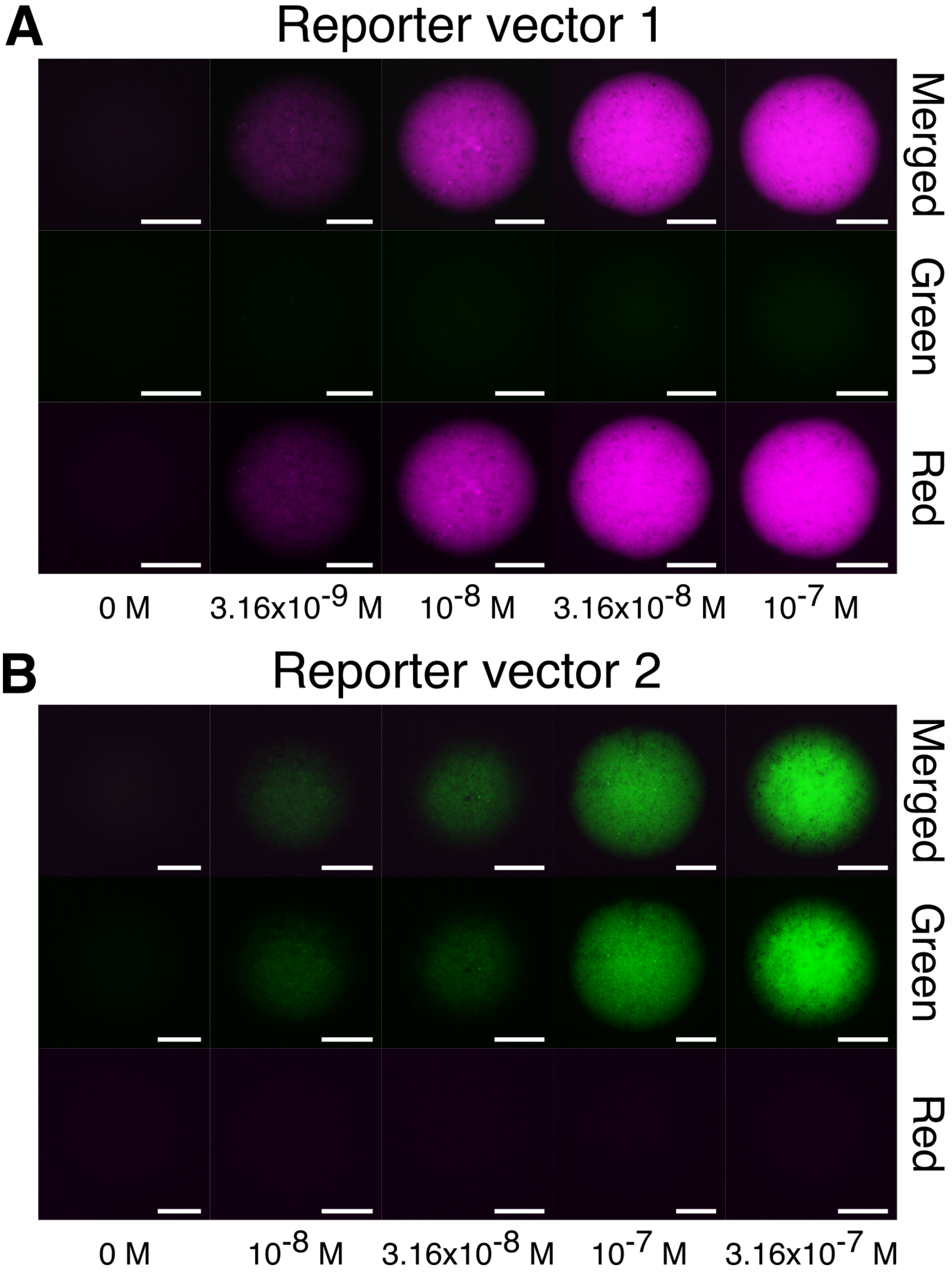

Figure S6: **Induction of red and green fluorescence by C6HSL in colonies of spherical cells carrying different versions of the reporter vector.** Red and green fluorescence of colonies generated from cells carrying the reporter vector 1 (A) and reporter vector 2 (B) grown on solid M9-glucose medium supplemented with different concentrations of C6HSL. Images were taken approximately 18 hours after inoculation. Scale bars 100 $\mu$m.

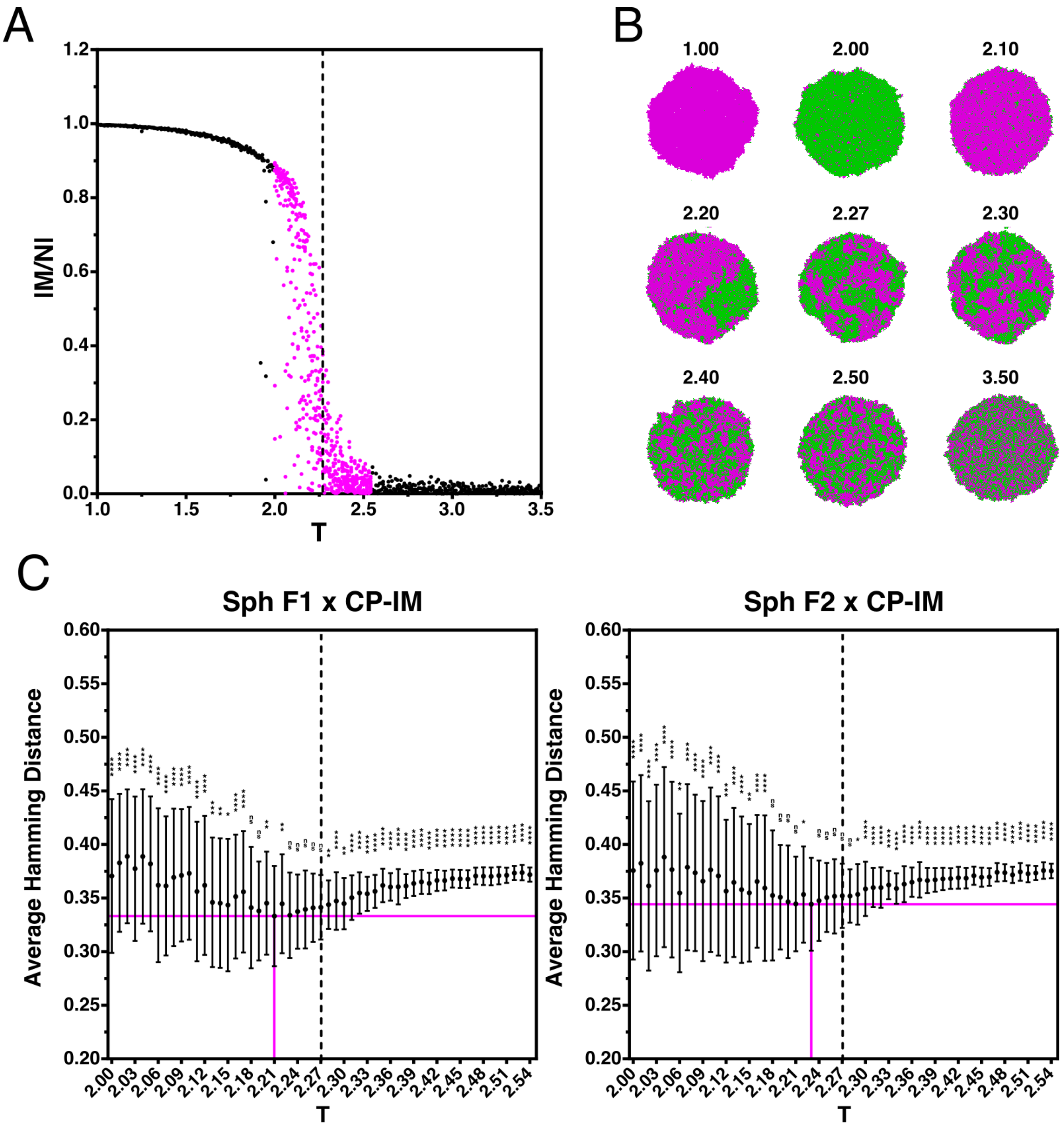

A
IM/NI
T
B
1.00
2.00
2.10
2.20
2.27
2.30
2.40
2.50
3.50
C
Sph F1 x CP-IM
Sph F2 x CP-IM
Average Hamming Distance
T

Figure S7: **Hamming distance between ferromagnetic colonies of spherical cells and simulated populations with CPIM.** (A) Absolute value of the magnetization per site $|M/N|$ as a function of $T$ for ferromagnetic populations simulated with CPIM. The dotted vertical line marks the critical value of $T$ for the Ising model ($T_c = 2.27$). Magenta dots correspond to the simulations used to calculate the Hamming distance in C. (B) Images of simulated ferromagnetic populations at different values of the control parameter $T$, highlighting that the probability of finding populations in the magenta or green state below $T_c$ is the same. (C) Hamming distance calculated between ferromagnetic colonies of spherical (Sph) cells carrying the reporter vector 1 (left, Sph F1) or 2 (right, Sph F2) and the simulated ferromagnetic populations with the CPIM model around the $T_c$. Black dots and error bars correspond to the mean $\pm$ the standard deviation of the Hamming distance calculated between the colonies (42 ferromagnetic colonies with reporter vector 1; 65 ferromagnetic colonies with reporter vector 2) and 10 simulated populations for $T$ between 2.00 and 2.54. For these simulations the following parameters were used: birth rate = 0.03, death rate = 0.00001, differentiation rate = 0.1, lattice = 256x256, R = 1 (nearest neighbors), and the images were captured on the 6800 generation. The value of the Hamming distance was divided by the total number of pixels. Dotted vertical lines mark the critical value of $T$ for the Ising model, and red solid lines mark the smallest average value of the Hamming distance. To compare the smallest average value of the Hamming distance with each other value, statistical analysis was performed using an ordinary One-Way ANOVA, followed by Dunnett's multiple comparisons test (95% confidence interval). ns (not significant): $P > 0.05$, $*$: $P \leq 0.05$, $**$: $P \leq 0.01$, $***$: $P \leq 0.001$, $****$: $P \leq 0.0001$.

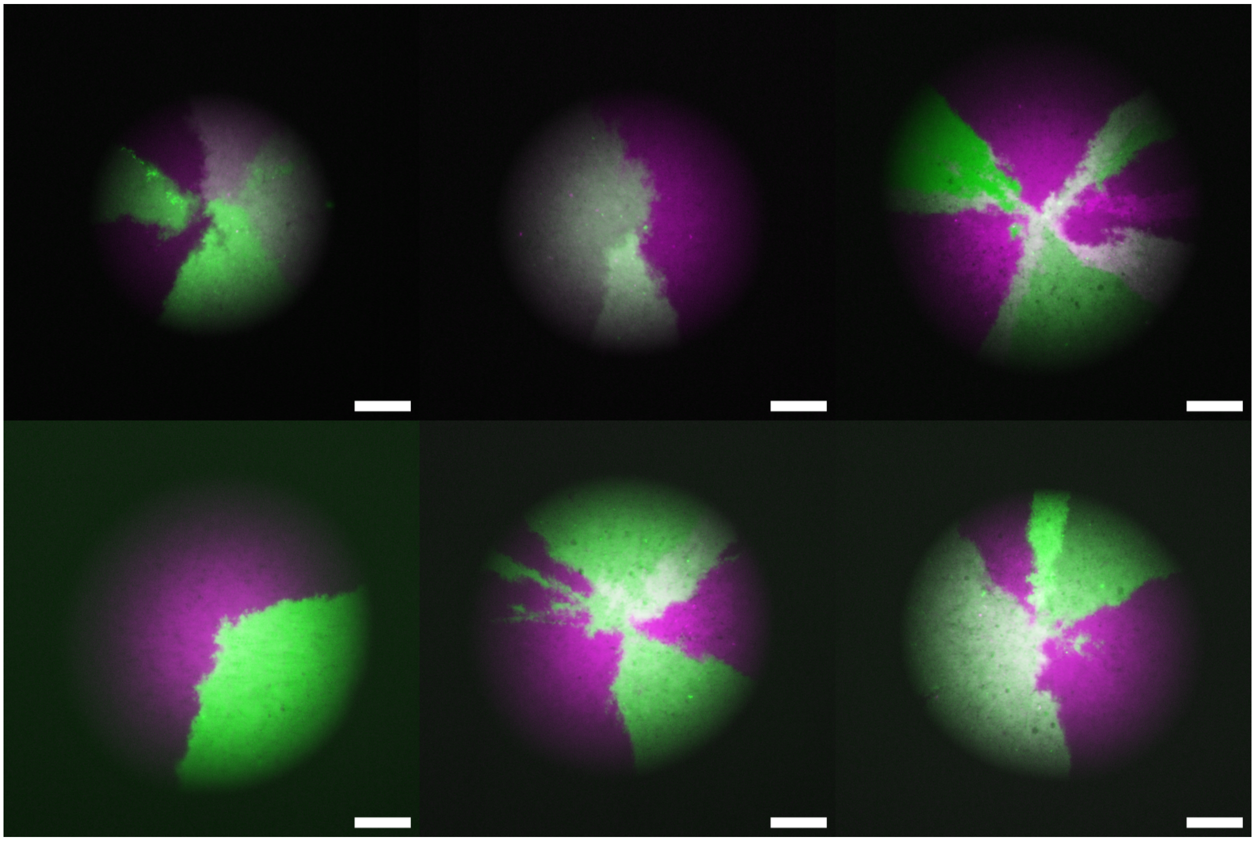

Figure S8: **Sharp boundaries generated between cellular state domains in colonies of spherical cells due to the segregation of fixed-state reporter vectors.** Representative images of the fluorescent patterns that emerge in colonies of spherical *E. coli* cells due to the segregation of fixed-state reporter vectors Nunez *et al* (2017). Each cell that gives rise to a colony contains two plasmids: one encoding a red fluorescent protein, and another encoding a green fluorescent protein. The patterns generated in these colonies are the result of the segregation of these plasmids during cell division: once a cell loses one of these plasmids, its cellular state and the cellular state of all its progeny is fixed, generating sharp boundaries between the cellular state domains. Scale bars 100 $\mu$m. The backbone plasmids used in this assay were obtained from Nunez *et al* (2017).

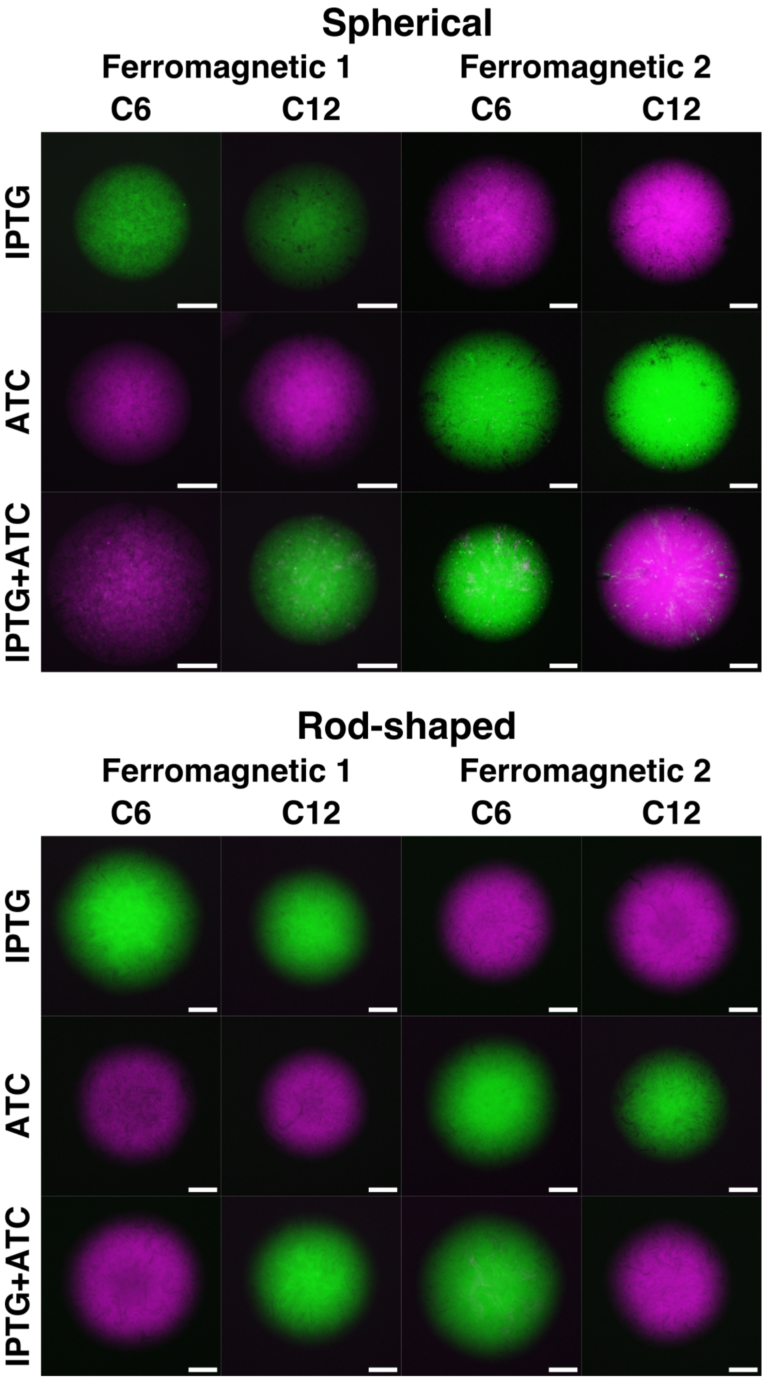

Figure S9: **Cellular state switch in ferromagnetic colonies grown on solid media supplemented with inhibitors.** Representative images of colonies of spherical (top) and rod-shaped (bottom) ferromagnetic cells that carry the reporter vector 1 (Ferromagnetic 1) or the reporter vector 2 (Ferromagnetic 2). Cells were grown on M9-glucose solid media supplemented C6HSL or C12HSL and with IPTG, ATC, or both inhibitors. These inhibitors bind to the repressors LacI and TetR, preventing them from repressing the expression of target genes. Thus, depending on the inhibitor present in the media, one state cannot be repressed, resulting in the repression of the other state, regardless of the coupling signal in which they are grown. Blocking both repressors at the same time, allows cells to adopt the cellular state induced by the coupling molecule present in the medium. Scale bars 100 $\mu$m.

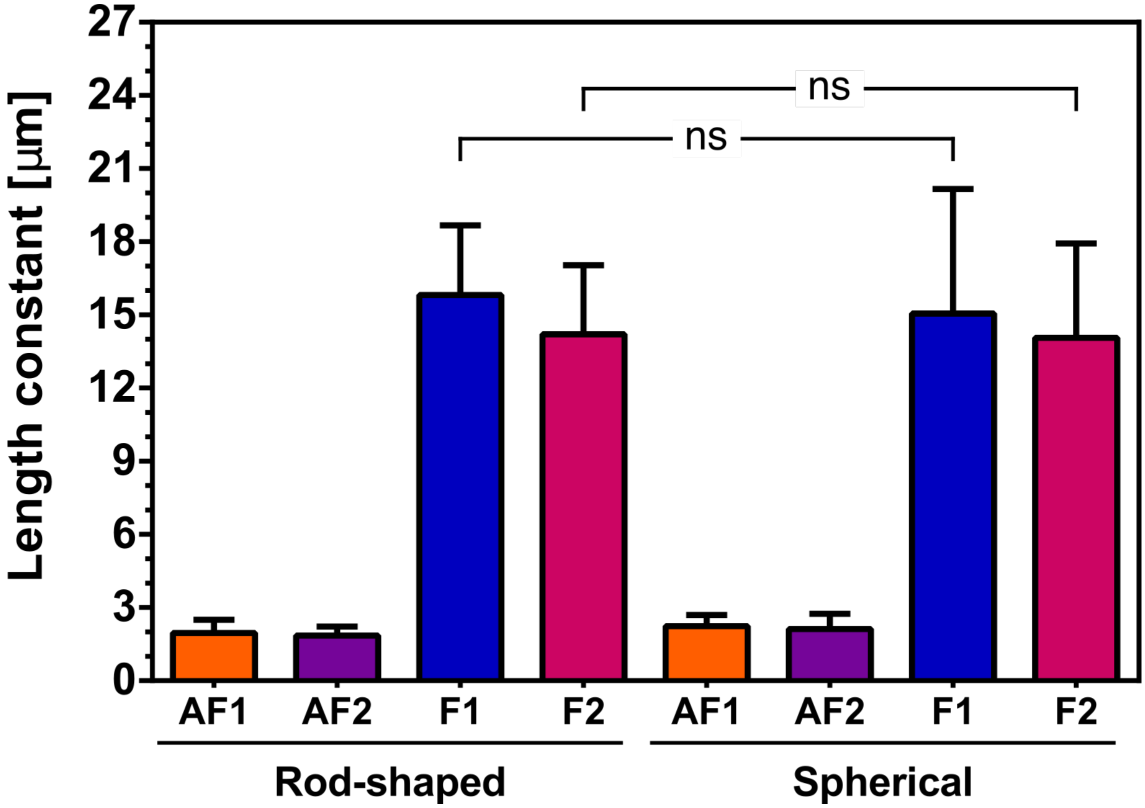

Figure S10: **Characteristic size of cellular states domains of ferromagnetic and anti-ferromagnetic colonies.** Length constants of ferromagnetic and anti-ferromagnetic colonies generated from rod-shaped and spherical cells, highlighting that there are no significant differences (P value: 0.0557 for F1 rod vs F1 sph and 0.4303 for F2 rod vs F2 sph) between the values of the length constants of ferromagnetic colonies of rod-shaped and spherical cells. Values correspond to the mean ± the standard deviation of the length constants of equation $y = y_0 * \exp(-x/b) + C$ to the data of sACF analysis, where the value of $b$ corresponds to the length constant. Around 40 colonies for each system were used. Statistical analysis was performed using an unpaired two-tailed Mann-Whitney test ($\alpha = 5\%$). ns (not significant): P $>$ 0.05.

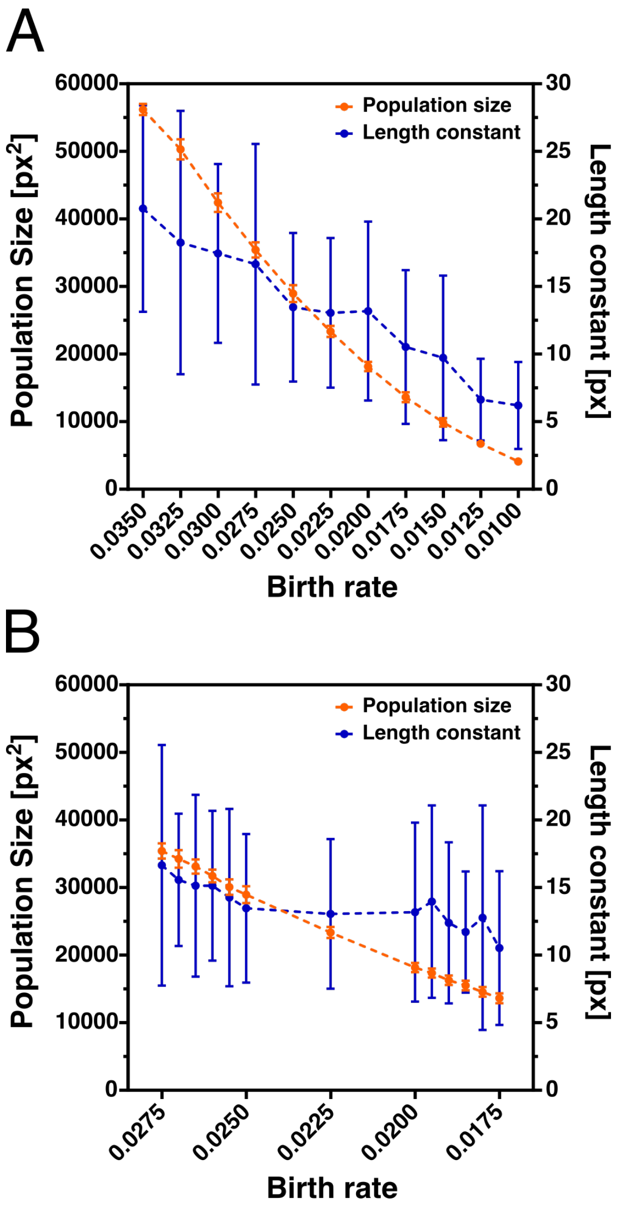

Figure S11: **Characteristic size of cellular state domains and the average size of simulated populations with ferromagnetic interactions.** A) Population area and length constant of ferromagnetic populations simulated with CPIM as a function of the birth rate. For these simulations the following parameters were used: death rate = 0.00001, differentiation rate = 0.1, lattice = 256x256, R = 1 (nearest neighbors), and the images were captured on the 6800 generation. While the size of the simulated populations decrease as the value of the birth rate decreases, the mean size of the cellular states domains (the length constant) remains constant between birth rate equal to 0.0250 and 0.0200 (P value: 0.4977 for 0.0225 vs 0.0250, 0.5131 for 0.0200 vs 0.0250). B) Adding more points to the analysis shows that there are no significant differences in the length constant between birth rates equal to 0.0255 and 0.0180 (P value: 0.8895 for 0.0250 vs 0.0255, 0.4107 for 0.0225 vs 0.0255, 0.2890 for 0.0200 vs 0.0255, 0.8436 for 0.0195 vs 0.0255, 0.1402 for 0.0190 vs 0.0255, 0.0502 for 0.0185 vs 0.0255, 0.0606 for 0.0180 vs 0.0255). Between these values of birth rate, the size of the population is reduced by more than half. Statistical analysis was performed using unpaired two-tailed Mann-Whitney test ($\alpha = 5\%$)
